# Redox-dependent condensation and cytoplasmic granulation by human ssDNA binding protein 1 delineate roles in oxidative stress response

**DOI:** 10.1101/2023.07.25.550517

**Authors:** Gábor M. Harami, János Pálinkás, Zoltán J. Kovács, Bálint Jezsó, Krisztián Tárnok, Hajnalka Harami-Papp, József Hegedüs, Lamiya Mahmudova, Nóra Kucsma, Szilárd Tóth, Gergely Szakács, Mihály Kovács

## Abstract

Human single-stranded DNA binding protein 1 (hSSB1/NABP2/OBFC2B) plays central roles in the repair of DNA breaks and oxidized DNA lesions. Here we show that hSSB1 undergoes liquid-liquid phase separation (LLPS) that is redox-dependent and requires the presence of single-stranded DNA or RNA, features that are distinct from those of LLPS by bacterial SSB. hSSB1 nucleoprotein droplets form under physiological ionic conditions, in response to treatment resulting in cellular oxidative stress. hSSB1’s intrinsically disordered region (IDR) is indispensable for LLPS, whereas all three cysteine residues of the oligonucleotide/oligosaccharide-binding (OB) fold are necessary to maintain redox-sensitive droplet formation. Proteins interacting with hSSB1 show selective enrichment inside hSSB1 droplets, suggesting tight content control and recruitment functions for the condensates. While these features appear instrumental for genome repair, we also detected hSSB1 condensates in the cytoplasm in response to oxidative stress in various cell lines. hSSB1 condensates colocalize with stress granules, implying unexplored extranuclear roles in cellular stress response. Our results suggest novel, condensation-linked roles for hSSB1, linking genome repair and cytoplasmic defense.

## INTRODUCTION

Single-stranded (ss) DNA binding (SSB) proteins are found in all living organisms (*1*). Their main function is the binding and stabilization of exposed ssDNA regions that form during DNA metabolic processes including replication, recombination, and repair (*2*). Protection of ssDNA regions from nucleolytic cleavage, unproductive secondary structure formation and harmful chemical alterations is essential for genome integrity. Moreover, SSBs specifically interact with several other proteins participating in genome maintenance, conferring an organizing hub function for SSBs in DNA metabolism (*1*, *3*). A 19-kDa subunit of the “prototypic” homotetrameric SSB of *Escherichia coli* (EcSSB) comprises an N-terminal oligonucleotide/oligosaccharide binding (OB) fold followed by an intrinsically disordered region (IDR) (*4*) (**Fig. 1A**). Besides mitochondrial SSB, the heterotrimeric replication protein A (RPA) was considered as the only functional metazoan SSB until the discovery of SSB homologs SSB1 and SSB2 (termed hSSB1/NABP2/OBFC2B and hSSB2/NABP1/OBFC2A, respectively, in humans) (*5*). hSSB1 and hSSB2 share high structural similarity to EcSSB, having an N-terminal OB fold followed by C-terminal IDR (**Fig. 1A-B**) (*4*, *6*). hSSB1 is expressed ubiquitously in human tissues, while hSSB2 expression is tissue specific, restricted mainly to immune cells and testes (*7*). Growing evidence highlights central functions for hSSB1 in DNA repair (*8*), telomere maintenance (*9*), RNA transcription (*10*), and embryonic development (*11*). While hSSB1 appears to perform wide-ranging cellular functions, most of our knowledge comes from studies focusing on its DNA repair function. hSSB1 was shown to be an early sensor of DNA damage upon ionizing radiation (IR), whereby it localizes to DNA double strand break (DSB) repair foci and organizes subsequent homologous recombination (HR) mediated DNA repair (*5*, *12*). hSSB1 facilitates DNA end resection, an early step in HR, by recruiting the MRN complex (*13*) and stimulating Exo1 activity (*14*). D-loop formation by RAD51 recombinase, a key step in early HR, is also facilitated by hSSB1 (*5*). hSSB1 colocalizes with Bloom’s syndrome helicase (BLM) on IR-induced DSBs (*15*). While hSSB1 is dispensable for normal S-phase DNA replication, it is required for stabilization and restart of replication forks that collapse upon hydroxyurea treatment (*16*). Another major role of hSSB1 is exerted during the repair of oxidative DNA damage (such as formation of 8- oxoguanine) via the hOGG1-mediated base excision repair (BER) pathway. hSSB1 relocalizes to chromatin after H_2_O_2_-induced oxidative stress to recruit hOGG1 to sites of damage and to stimulate its incision activity (*8*).

**Fig. 1.**
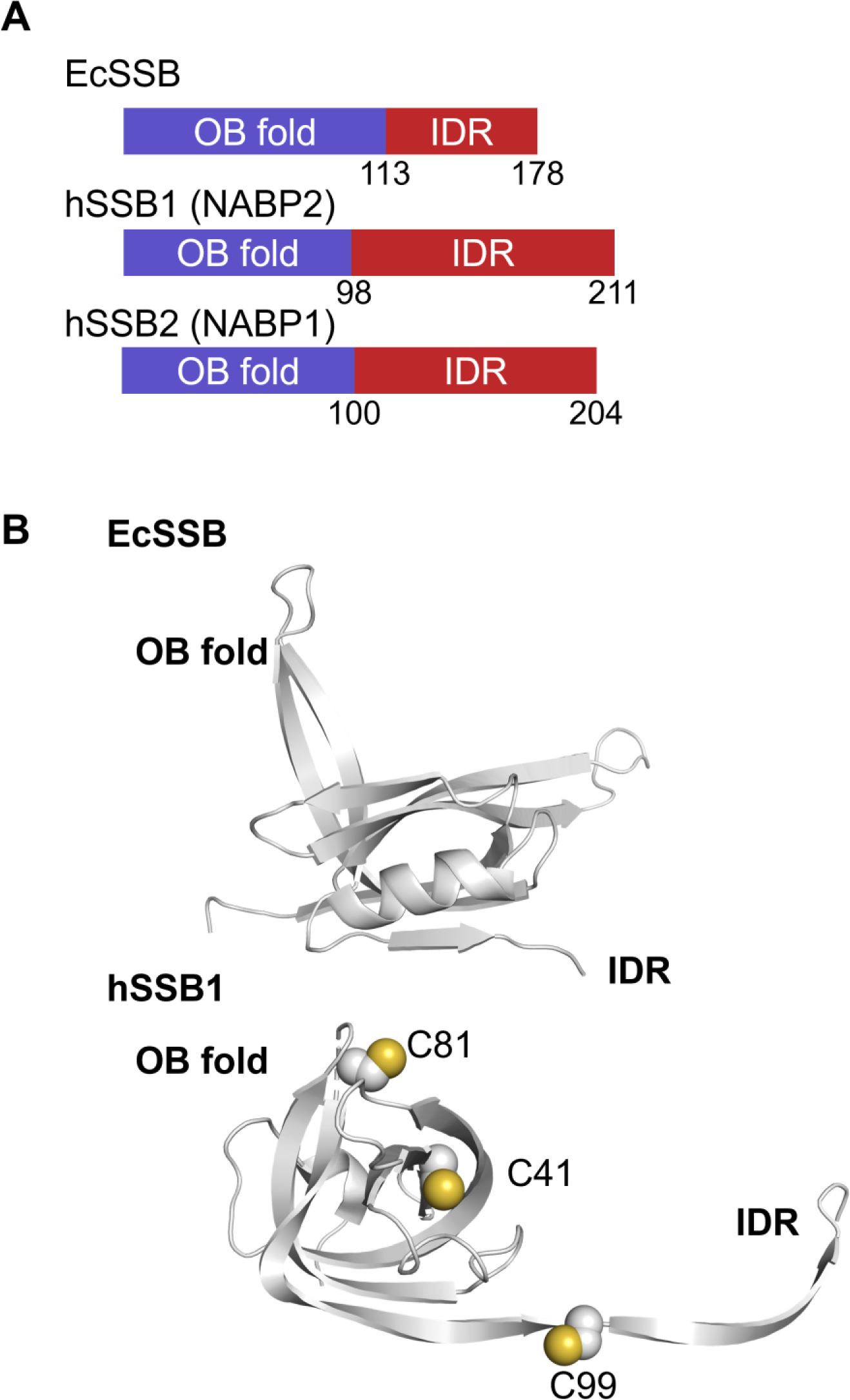
Bacterial and human SSB proteins show high structural similarity. (**A**) Domain structure of *E. coli* (Ec) and human (h) SSB proteins, with amino acid positions indicated at domain boundaries. (**B**) Three-dimensional structures of EcSSB (PDB code 4MZ9) and hSSB1 (5D8F), with cysteine residues highlighted. (EcSSB contains no cysteines.) Structures shown are visible until aa 114 and 110, respectively.

While EcSSB is a stable homotetramer, the ability of hSSB proteins to self-oligomerize appears more dynamic and is of functional significance. Under reducing conditions, hSSB1 was shown to be dominantly monomeric (*6*, *17*). However, unlike EcSSB that lacks cysteines, hSSB1 contains three conserved cysteine residues (at amino acid (aa) positions 41, 81, and 99; **Fig. 1B**) that could potentially facilitate redox-dependent covalent oligomerization via disulfide bonds. hSSB1 was indeed shown to oligomerize covalently upon oxidative stress, with this feature being abolished by the C99S or C41S substitutions (*17*).

Based on the observed oligomerization propensity and tetrameric structure of other SSBs, Touma et al. proposed a structural model in which the hSSB1 tetramer is stabilized by C81- C81 and C99-C99 intermolecular disulfide bridges, whereas C41 buried inside the OB fold may act as a redox sensor, with its oxidative status allosterically influencing protein structure (*18*). Although the structures of hSSB1 oligomeric forms have not been experimentally clarified, oxidation-dependent hSSB1 oligomerization was found necessary for efficient hOGG1-mediated BER and also for enhanced hSSB1 binding to DNA containing oxidative lesions (*8*), but appears dispensable for DSB repair (*17*). Furthermore, hSSB1 was observed as a monomer in the ternary SOSS (sensor of ssDNA) complex formed with Integrator complex subunit 3 (INTS3) and SOSS complex subunit C (INIP, C9Orf80, hSSBIP1) proteins (*19*, *20*). hSSB1 directly interacts with INTS3, and the complex plays an essential role in HR-mediated DSB repair via ATM activation, RAD51 recruitment to DNA damage (*20–22*), and facilitation of DSB resection by Exo1 (*14*). Complex formation is independent of DNA damage, while INTS3 knockdown results in a decrease in nuclear hSSB1 foci after irradiation. INTS3 is required for the transcription of hSSB1; thus, the knockdown phenotype is rescued by ectopic expression of hSSB1 (*21*). hSSB1 oligomerization proposedly does not interfere with INTS3 interaction (*18*). Taken together, these data point to the functional importance of various homo- and heterooligomeric forms of hSSB1 in stress response.

We recently discovered that EcSSB forms liquid-like dynamic condensates under physiological conditions *via* liquid-liquid phase separation (LLPS) (*23*). SSB-interacting proteins are selectively enriched inside droplets, while droplet formation is inhibited by ssDNA. Together with earlier data on subcellular distribution of EcSSB (*24*), these findings led to a model in which EcSSB condensates serve as organizers of bacterial genome maintenance (*23*). Upon genomic stress resulting in exposed ssDNA, droplet contents (EcSSB and partner proteins) can be rapidly deployed at the site of action. In this work we also raised the possibility of hSSB1 (and hSSB2) condensation based on *in silico* predictions (*23*).

Here we show that hSSB1 indeed forms LLPS condensates, with properties that markedly differ from those of bacterial SSB. hSSB1 droplet formation requires coacervation with nucleic acids and is tightly regulated by redox conditions, suggesting the importance of LLPS upon oxidative stress, e.g., in the repair of oxidative DNA lesions and/or in stress-related transcription regulation. Covalent oligomerization is not a prerequisite for hSSB1 LLPS; however, all hSSB1 cysteines are necessary for redox-dependent condensation. We also show that cytoplasmic hSSB1 droplets form in various cancerous and non-cancerous cell lines in response to oxidative stress, colocalizing with stress granules, pointing to unexplored extranuclear stress response roles for hSSB1. These results extend the emerging concept of nucleoprotein LLPS centrally contributing to genome maintenance and other areas of cellular stress response.

## RESULTS

### hSSB1-ssDNA and hSSB1-ssRNA nucleoprotein coacervates form selectively and reversibly under oxidative conditions

In line with our previous *in silico* predictions (*23*), epifluorescence microscopy experiments showed that purified hSSB1 is indeed able to undergo LLPS (**Figs. 2A, S1-2**). Importantly, however, unlike in the case of EcSSB, where ssDNA inhibited droplet formation (*23*), hSSB1 requires the presence of ssDNA or ssRNA for LLPS. In addition, we found that hSSB1 requires oxidative conditions for robust droplet formation (**Fig. 2A**), in line with the established role of the protein in oxidative DNA damage repair (*8*). Hydrogen peroxide (H_2_O_2_) is a common reactive oxygen species (ROS) *in vivo* (*25*), and it is an established agent to mimic oxidative stress conditions both *in vitro* and in cell culture experiments. Intriguingly, ssRNA induced hSSB1 droplet formation to a limited extent even in reducing conditions; however, LLPS induction by H_2_O_2_ was apparent even in this case (**Fig. 2A**). We found that hSSB1 binds ssDNA and ssRNA with similar affinities (**Fig. 2B, Table S1**); thus, the different LLPS propensities of the hSSB1-ssDNA and hSSB1-ssRNA nucleoproteins do not result from differences in the strength of protein-nucleic acid interactions. Liquid-like behavior of hSSB1 condensates is supported by spherical morphology and volume-additive fusion of droplets (**Fig. 2C**). In epifluorescence microscopy experiments, condensate formation became apparent at H_2_O_2_ concentrations in the micromolar range (**Fig. 2D**).

**Fig. 2.**
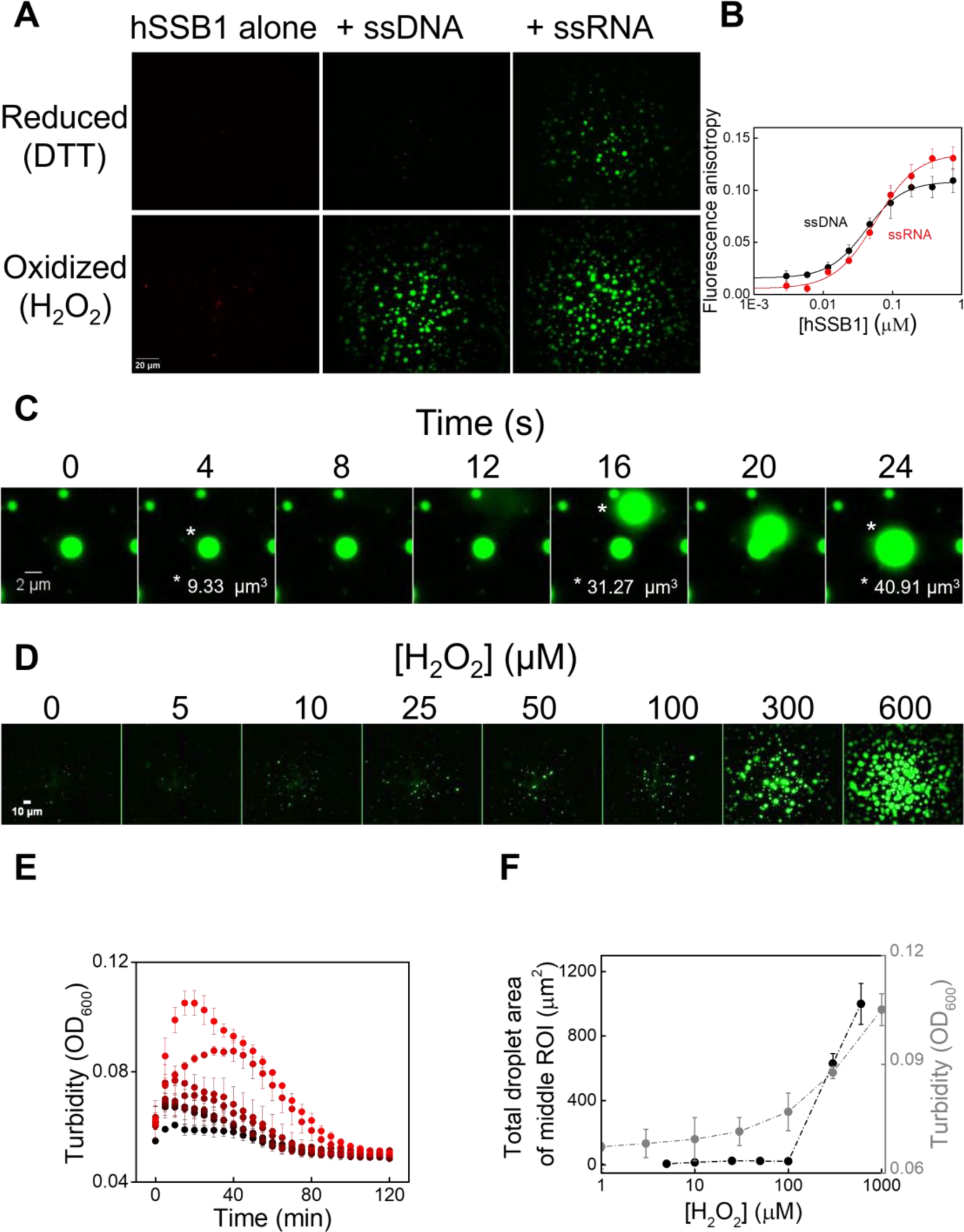
hSSB1 forms ssDNA and ssRNA nucleoprotein coacervates in a redox-dependent manner. (**A**) ssDNA and ssRNA induce hSSB1 LLPS in oxidative conditions. Representative epifluorescence microscopy images (*n* = 3 independent measurements) of hSSB1 (5 µM monomer concentration) were obtained in the absence of nucleic acids using AlexaFluor647- labeled hSSB1 (0.1 µM), and in the presence of ssDNA (2 μM dT_45_ containing 0.1 μM Cy3- dT_45_) and ssRNA (2 μM U_41_ containing 41-mer Cy3-labeled nonhomopolymeric ssRNA) (cf. **Fig. S2**) in reducing (1 mM DTT) and oxidizing (980 μM H_2_O_2_) conditions. (**B**) ssDNA and ssRNA (10 nM 3’-fluorescein labeled nonhomopolymeric 36-mer) binding by hSSB1 measured with fluorescence anisotropy titrations. Solid lines show fits using the Hill equation. Means ± SEM are shown for *n* = 3 independent experiments. Determined equilibrium dissociation constants (*K*_d_) and Hill-coefficients (*n*) are shown in **Table S1**. (**C**) Droplet fusion, volume additivity, and spherical morphology support liquid-like behavior of hSSB1 droplets. Time-lapse fluorescence images of fusion of hSSB1 droplets (5 μM) are shown, recorded in the presence of labeled ssDNA (2 μM dT_79_ containing 0.1 μM Cy3-dT_79_) and H_2_O_2_ (980 µM). (**D**) H_2_O_2_ concentration dependence of the appearance of hSSB1 droplets (5 µM hSSB1, 2 µM dT_79_ containing 0.1 µM Cy3-dT_79_). (**E**) Time-dependent turbidity measurements showing hSSB1 LLPS even at low micromolar H_2_O_2_ concentrations (2 µM hSSB1, 1 µM dT_79_ ssDNA; H_2_O_2_ concentrations from bottom to top were 0, 1, 3, 10, 30, 100, 300, and 1000 µM). Error bars represent SEM for *n* = 3 independent experiments. (**F**) H_2_O_2_ concentration dependence of total droplet area of middle ROIs in microscopy images (panel **D**) and maximal OD values from panel **E**. Error bars represent SEM for *n* = 3 independent measurements.

Turbidity measurements, which are sensitive to the appearance of small-sized droplets that are yet undetectable in the epifluorescence microscope, indicated phase separation at H_2_O_2_ concentrations in the low micromolar range, a physiologically relevant regime for cellular oxidative stress (*25*) (**Fig. 2E-F**). Turbidity and microscopy measurements indicated rapid appearance of condensates (< 10 minutes) in response to H_2_O_2_ treatment, with turbidity eventually decreasing over longer time scales due to droplet fusion and settling, phenomena also seen previously with EcSSB (*23*) (**Figs. 2C, E; S3A**). Importantly, EcSSB shows LLPS independent of redox conditions, underscoring the unique nature of redox-sensitive LLPS by hSSB1 (**Fig. S3B**). Furthermore, hSSB1 droplet formation was partially reversible upon addition of the reducing agent DTT (**Fig. S3C-D**). Prominent droplets disappeared after 1h incubation, leaving only a faint signal on the microscope slide surface. As LLPS systems are generally sensitive to the ionic milieu, we also tested the effect of various salts on hSSB1 LLPS. High, supraphysiological salt concentrations inhibited droplet formation (**Fig. S4**), and addition of KCl at supraphysiological concentration rapidly dissolved droplets after 5 seconds (**Fig. S5**). Importantly, however, in a phosphate-buffered (pH 7.5), quasi-physiological milieu of 140 mM K^+^, 10 mM Na^+^, 10 mM Cl^-^, 8 mM magnesium acetate, and 30 mg/ml PEG_20000_ to mimic crowding, hSSB1 exhibited LLPS in the presence of ssDNA in response to H_2_O_2_ treatment (**Fig. S4E**). Taken together, these findings reflect the propensity of hSSB1 for redox-dependent nucleoprotein coacervate formation under physiological intracellular chemical conditions.

### hSSB1 LLPS shows bell-shaped dependence on nucleic acid concentration

In addition to the factors explored above, the total protein concentration and the protein-DNA stoichiometry could also influence LLPS behavior. Thus, we investigated the dependency of phase separation on hSSB1 and ssDNA concentrations. Using epifluorescence microscopy, we found that in the presence of a fixed amount of ssDNA, large phase separated droplets started to appear when the bulk hSSB1 concentration reached a threshold around 1 μM (hSSB1 monomer concentrations are stated throughout this article unless otherwise indicated) (**Fig. 3A**). Turbidity measurements also confirmed this observation (**Fig. 3B**), and protein concentration-dependent droplet formation was similar in the presence of ssDNA and ssRNA (**Fig. S6A**). Using a fixed amount of hSSB1 we found that the formation of phase-separated droplets showed a bell-shaped dependence on both ssDNA and ssRNA concentration (**Figs. 3C-D, S6B**).

**Fig. 3.**
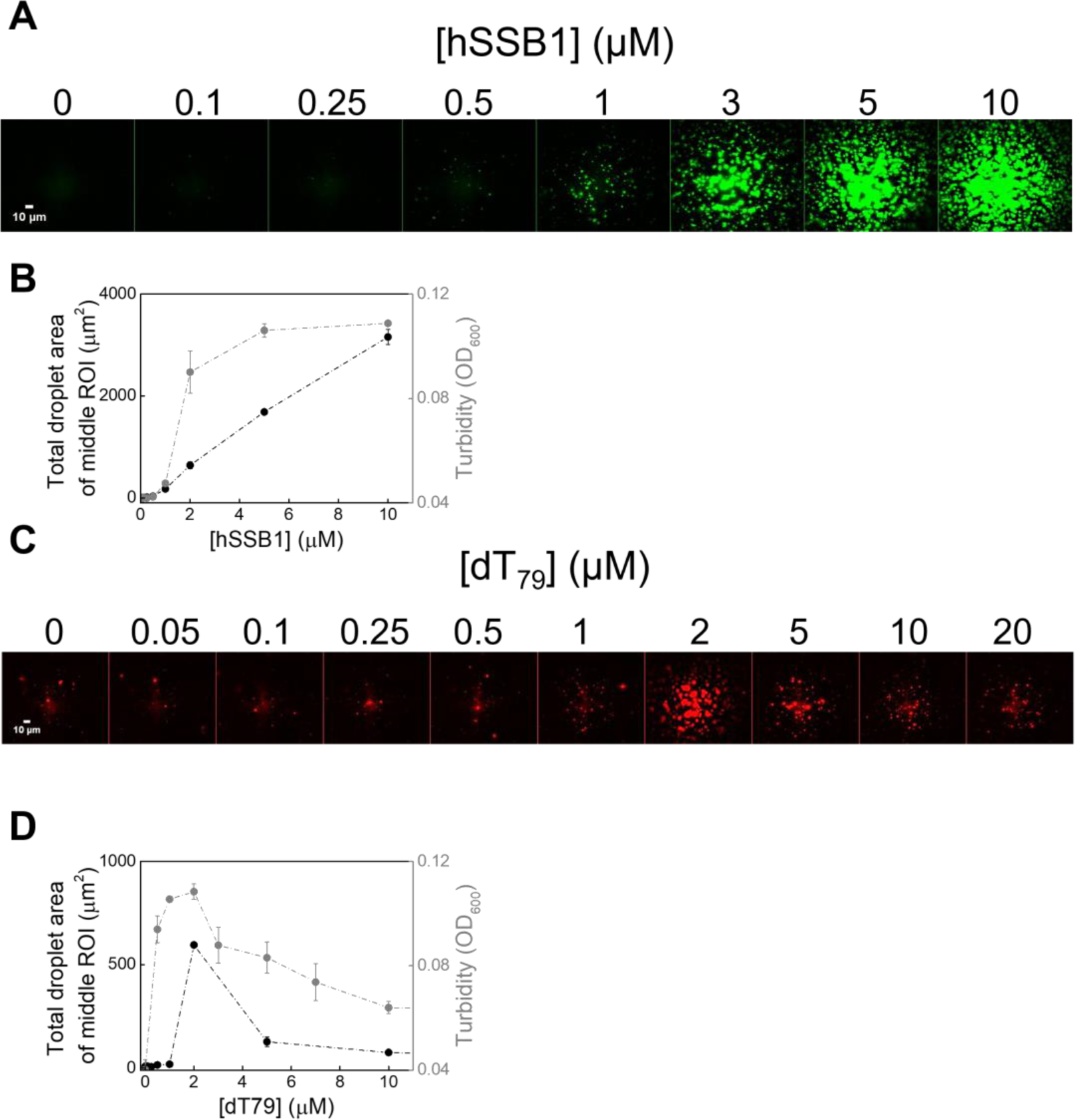
hSSB1 nucleoprotein condensation occurs in a stoichiometric fashion, with a bell- shaped dependence on nucleic acid concentration. (**A**) Epifluorescence microscopy images (*n* = 3 independent measurements) recorded in the presence of labeled ssDNA (2 µM dT_79_ containing 0.1 µM Cy3-dT_79_) and H_2_O_2_ (980 μM) showing that hSSB1 droplets become apparent at 1 μM protein concentration. (**B**) hSSB1 concentration dependence of total droplet area of middle ROIs in microscopic images (panel **A**) and OD_600_ values from turbidity measurements in the presence of ssDNA (2 μM dT_79_) and H_2_O_2_ (980 μM). Error bars represent SEM for *n* = 3 independent measurements. (**C**) Epifluorescence microscopic images (*n* = 3 independent measurements) of fluorescently labeled hSSB1 droplets (5 µM hSSB1 containing 0.1 µM Alexa Fluor647-labeled hSSB1) titrated with dT_79_ in the presence of 980 µM H_2_O_2_. (**D**) dT_79_ concentration dependence of total droplet areas from fluorescence microscopy experiments shown in panel **C** and OD_600_ values from turbidity measurements using 5 μM hSSB1 in the presence of 980 μM H_2_O_2_. Error bars represent SEM for *n* = 3 independent measurements.

In addition to the above experiments, we tested the dependence of droplet formation on nucleic acid length. We measured turbidity of hSSB1 samples in the presence of dT homopolymers ranging in size from 18 to 96 nucleotides (nt) (**Fig. S6C**). To avoid possible inhibitory effects of salts on LLPS and/or ssDNA binding, measurements were carried out in no-salt LLPS buffer (see Methods). dT_18_ was unable to trigger LLPS, while longer dT species showed bell-shaped phase diagrams with maximal turbidity values corresponding to an ssDNA interaction stoichiometry of around 10-15 nt ssDNA / hSSB1 monomer. This finding may indicate architectural differences between hSSB1 and EcSSB nucleoproteins, with the latter binding about 35 nt of oligonucleotide by two subunits of a tetramer in LLPS-competent conditions (*23*). Based on a binding site size of 10-15 nt and on the observation that dT_18_ was unable to promote LLPS, the data imply that at least two hSSB1 monomers must interact with an ssDNA molecule in order to undergo phase transition.

### The intrinsically disordered region (IDR) is required for strong nucleic acid binding and LLPS

Previously we found that the IDR of EcSSB is essential for LLPS: IDR-truncated EcSSB showed no LLPS propensity and formed amorphous aggregates (*23*). In addition, the amino acid composition of the EcSSB IDR has also been shown to be an important determinant of LLPS (*26*). Since hSSB1 also contains a long (113-aa) IDR (**Figs. 1, 4A**), and our previous *in silico* analysis predicted high LLPS propensity for this region (*23*), we tested the LLPS propensity of an hSSB1 variant lacking the IDR region (hSSB1-dIDR). We observed no droplet formation for this construct even in the presence of ssDNA and H_2_O_2_; only aggregates were observed at higher protein concentrations (**Fig. 4A**). We also found that hSSB1-dIDR shows an about 10-fold decrease in ssDNA binding affinity, compared to the wild-type protein (**Fig. 4B, Table S1**); however, hSSB1-dIDR droplets did not form even at quasi-saturating ssDNA concentrations in the presence of H_2_O_2_ (**Fig. S7**). These data demonstrate that the IDR has a moderate contribution to ssDNA binding, but is indispensable for phase separation.

**Fig. 4.**
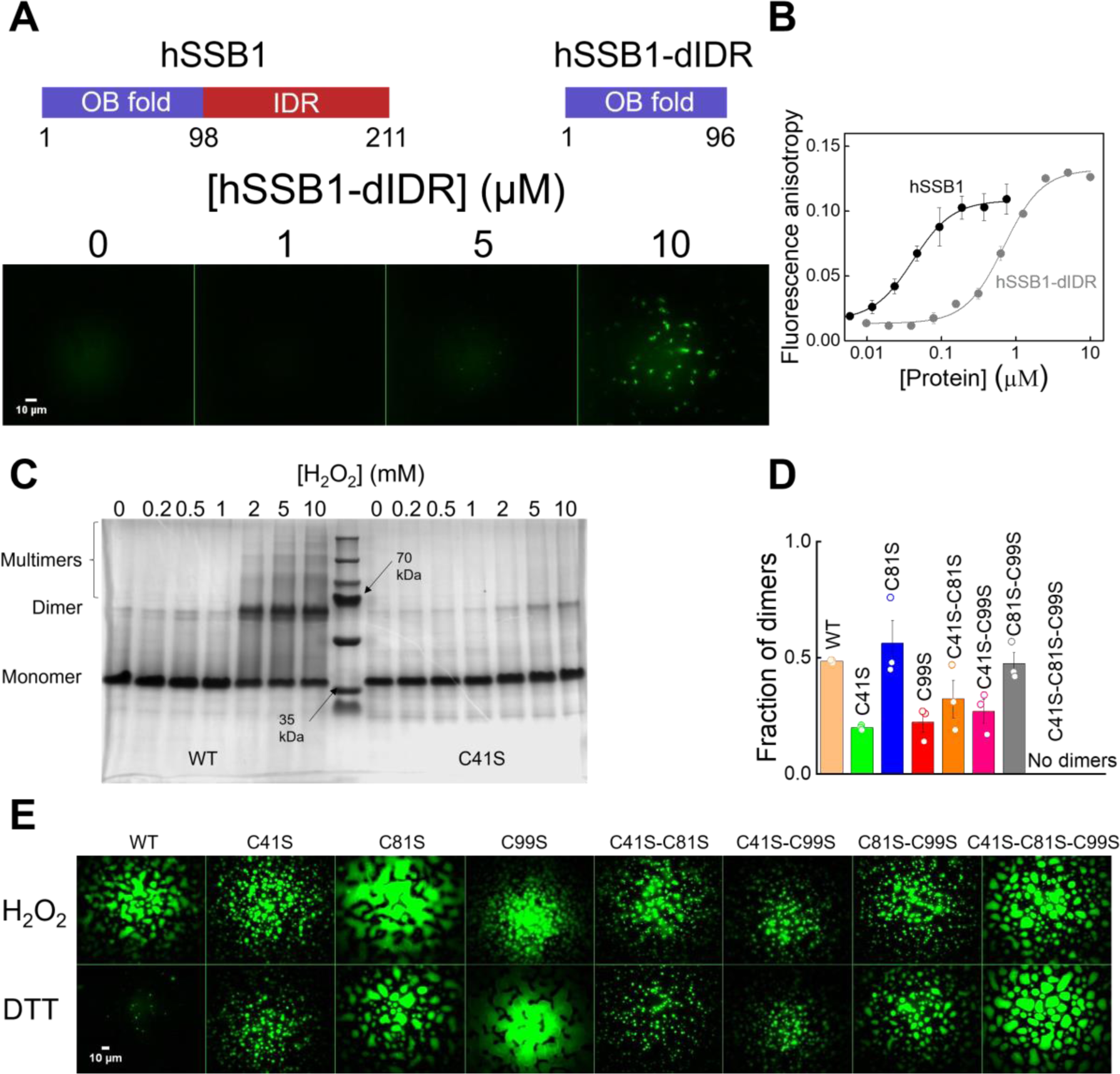
Intrinsically disordered region. (**IDR**) **of hSSB1 is indispensable for LLPS propensity, cysteines 41 and 99 mediate covalent oligomerization, while all hSSB1 cysteines are required for redox-sensitive condensation.** (**A**) Domain structure of hSSB1 WT and hSSB1-dIDR constructs (top panel). Epifluorescence images (*n* = 3 independent measurements) of hSSB1-dIDR in the presence of labeled ssDNA (2 µM dT_79_ containing 0.1 µM Cy3-dT_79_) and H_2_O_2_ (980 µM) show no droplet formation, only amorphous aggregates at high protein concentration (bottom panel). (**B**) ssDNA (10 nM 3’-fluorescein labeled nonhomopolymeric 36-mer) binding of hSSB1 and hSSB1-IDR measured in fluorescence anisotropy titrations show a marked decrease in the ssDNA affinity of hSSB1-dIDR compared to hSSB1 WT. Solid lines show fits using the Hill equation. Means ± SEM are shown for *n* = 3 independent measurements. Determined equilibrium dissociation constants (*K*_d_) and Hill coefficients (*n*) are shown in **Table S1**. (**C**) A representative electrophoretogram (*n* = 3 independent measurements) of WT and C41S hSSB1 variants shows changes in monomer : dimer ratio in response to H_2_O_2_ treatment. Densitometric analysis was applied to determine relative protein amounts in individual fractions (cf. **Figs. S1 and S8**). Monomers and dimers separated clearly in every case, while higher-order structures were distinguishable only in some cases, and thus omitted from the analysis. Note that the electrophoretic mobility of monomeric hSSB1 is lower than expected based on its molecular weight of 22 kDa, which is a frequently observed feature for proteins containing long ID regions. Please note that each oxidation reaction contained < 100 μM DTT reducing agent originating from the storage buffer of hSSB1 constructs. (**D**) Fraction of dimers formed by hSSB1 variants in response to the highest applied H_2_O_2_ concentration (10 mM) (cf. **Fig. S8**). The data indicate the key contributions of C41 and C99 to covalent dimerization (see Results). Means ± SEM are shown together with individual data points from *n* = 3 independent measurements. (**E**) Epifluorescence microscopy images (*n* = 3 independent measurements) showing that all hSSB1 variants retained their ability to undergo LLPS, but the redox sensitivity was lost for all non-WT cysteine variants (5 µM protein, 2 µM dT_79_ containing 0.1 µM Cy3-dT_79_, 980 µM H_2_O_2_ or 1 mM DTT were present in all samples).

### C41 and C99 are required for redox-dependent covalent oligomerization

A possible mechanism underlying the observed redox-sensitive condensation (**Figs. 2A, S3A**) is covalent oligomerization of hSSB1 through (some of) its three conserved cysteine residues (C41, C81, C99) (**Fig. 1B**), which could increase the multivalency of protein-protein and protein-nucleic acid interactions needed for LLPS (*27*, *28*). C41 was shown to be important for redox-dependent covalent oligomerization, but since it is not exposed to solvent, it was hypothesized to allosterically influence the formation of disulfide bridges between C81 and C99 residues *via* its oxidative state (*18*). To determine the contribution of individual hSSB1 cysteines (**Fig. 1B**) to covalent self-oligomerization and LLPS, we generated all possible Cys-to-Ser substitution variants (C41S, C81S, C99S, C41S-C81S, C41S-C99S, C81S-C99S, and C41S-C81S-C99S). We tested the propensity of hSSB1 variants for covalent oligomerization upon H_2_O_2_ treatment *via* SDS-PAGE densitometry (**Figs. 4C-D, S8**). We found that covalent dimers were the major oligomeric species formed upon oxidation, with discernible appearance of higher-order oligomers in some cases (**Fig. S8A**). Upon H_2_O_2_ treatment, about half of the total wild type (WT) hSSB1 pool formed dimers at saturating H_2_O_2_ concentration (**Figs. 4D, S8B**). C41S and C99S variants showed reduced dimer formation, while C81S retained the WT phenotype. For the C41S-C81S and C41S-C99S double substituted variants, we observed similar inhibition of dimerization to that for C41S and C99S. Interestingly, C81S-C99S rescued the WT phenotype, and its initial distribution in the absence of H_2_O_2_ showed a higher dimer : monomer ratio than that for WT hSSB1, implying that the combined absence of C81 and C99 thiols rendered C41 capable of disulfide formation (**Fig. S8B**). As expected, the triple substituted variant C41S-C81S-C99S showed no covalent oligomerization. We note that all hSSB1 variants, except hSSB1-dIDR, contained a minor non-covalent dimeric species corresponding to 70 kDa (cf. **Figs. S1, S8**), which was resistant to SDS and DTT treatment, but did not affect the determination of the extent of oxidation-induced hSSB1 covalent oligomerization. Taken together, these data show that both C41 and C99, but not C81, crucially contribute to redox-dependent covalent dimerization.

### All cysteines of hSSB1 are required for redox-regulated LLPS propensity

After asserting that the ssDNA binding properties of the above described hSSB1 variants were not affected by the amino acid substitutions (**Fig. S9A, Table S1**), we also investigated the LLPS propensity of these hSSB1 variants (**Fig. 4E**). Interestingly, all constructs retained their ability to undergo LLPS, but the redox sensitivity of this feature was lost in all substituted variants: phase-separated droplets were visible both in the presence of H_2_O_2_ and in DTT. However, the dependence of droplet formation on the presence of ssDNA was unaffected; variants were unable to undergo LLPS in the absence of ssDNA (**Fig. S9B**). In the presence of ssDNA, C41S and C99S, which were less efficient in covalent oligomerization, and even C41S-C81S-C99S showing no covalent oligomerization, were still able to form phase-separated droplets (**Fig. 4E**).

To ensure that the loss of redox sensitivity is not a result of inhomogeneity in starting materials, we performed LLPS measurements using freshly reduced WT and C41S-C81S- C99S hSSB1 samples following overnight incubation with TCEP reducing agent, followed by buffer exchange (**Fig. S10A**). We observed the same LLPS propensities as in the earlier experiments: the triple substituted variant remained insensitive to redox conditions, while the WT protein showed robust droplet formation only under oxidizing conditions. These findings showed that disulfide formation and covalent oligomerization *per se* are not needed for hSSB1 phase separation, but the presence of all cysteines is needed for redox-sensitive behavior.

To test whether H_2_O_2_ exerts its effect directly through cysteines, we also used the thiol-specific oxidant diamide instead of H_2_O_2_ (**Fig. S10B**). WT hSSB1 showed similar droplet formation in response to diamide treatment as it did in H_2_O_2_, highlighting the role of cysteine oxidation as a regulator of redox-sensitive hSSB1 LLPS. The redox-insensitive LLPS behavior of the C41S-C81S-C99S construct was unaffected by diamide treatment.

### Genome maintenance proteins are selectively enriched in hSSB1 droplets that act as molecular filters

hSSB1 was shown to promote recruitment of Bloom’s syndrome (BLM) helicase to double-stranded DNA breaks during HR (*15*). In addition, integrator subunit 3 (INTS3) is required for the transcription of hSSB1 (*21*) and serves as a major interacting partner of hSSB1 (and hSSB2) in the ternary SOSS complex, which is crucial for DSB repair (*20–22*) (**Fig. 5A**).

**Fig. 5.**
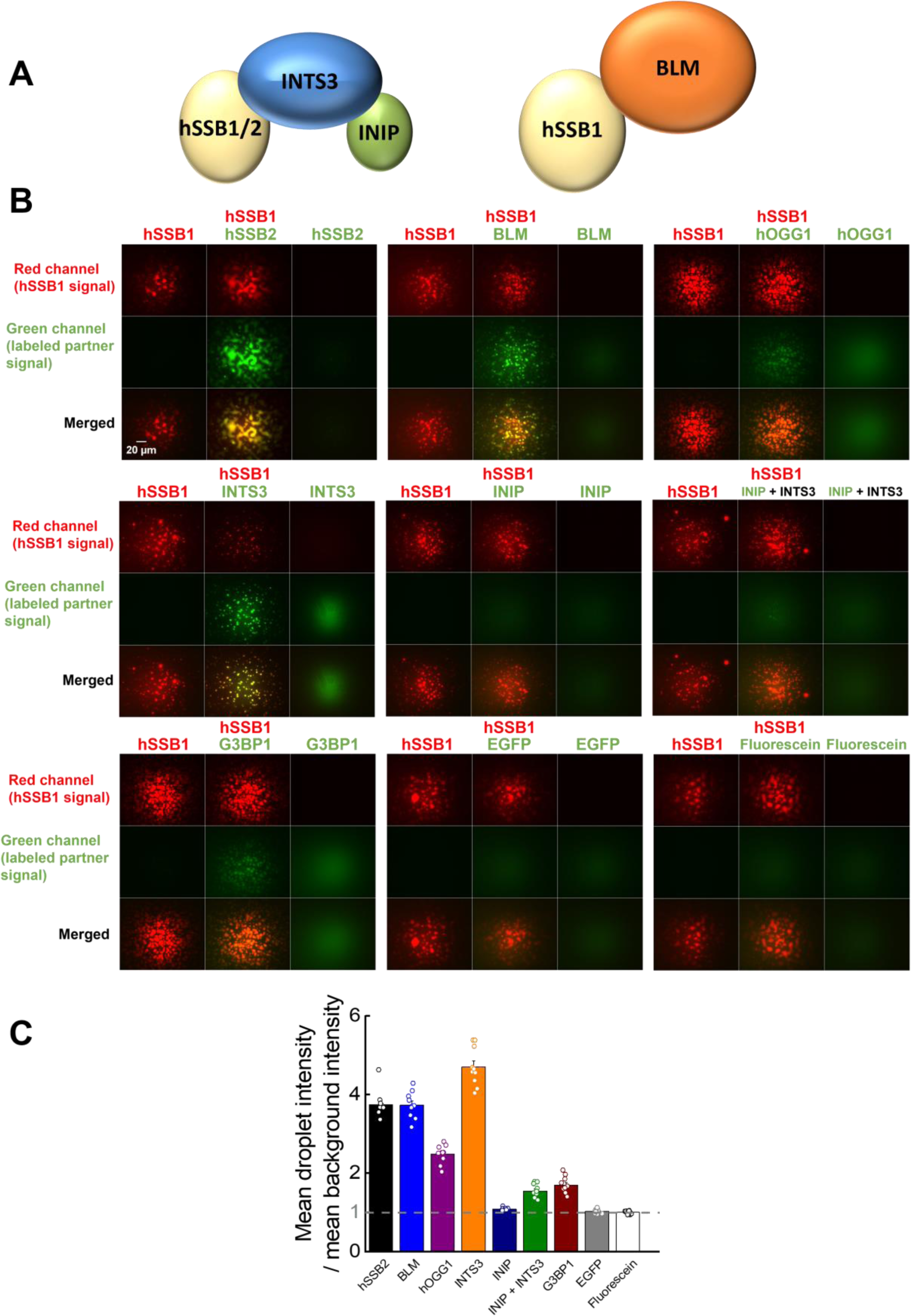
Genome maintenance proteins are selectively enriched inside hSSB1 droplets, reflecting interaction-based control of their content. (**A**) Schematic illustration of SOSS and hSSB1-BLM helicase complexes. INTS3 interacts both with INIP and hSSB1, while there is no direct interaction between INIP and hSSB1. hSSB2 can replace hSSB1 in the SOSS complex. (**B**) Two-channel fluorescence microscopy images (*n = 3* independent measurements) showing enrichment of proteins inside hSSB1 droplets. Columns represent three separate experiments (hSSB1, hSSB1 + partner, partner alone), while rows represent fluorescence channels. 5 µM hSSB1 with 0.1 µM AlexaFluor647-labeled hSSB1 were present in the samples, together with ssDNA (2 µM dT_79_) and H_2_O_2_ (980 µM). 180 nM labeled interaction partner was used. In experiments containing labeled INTS3, labeled ssDNA (2 µM dT_79_ containing 0.1 µM Cy3-dT_79_) was used to visualize hSSB1 droplets (5 µM unlabeled protein). See Materials and Methods for further details. Red channel shows hSSB1 droplets, green channel shows labeled interaction partners. Yellow color indicates co-condensation. Images were not background corrected. (**C**) Enrichment of various components in hSSB1 droplets, calculated as the ratio of the mean signal intensity within droplets and the mean background intensity, determined from background-uncorrected fluorescence images recorded for the indicated fluorescent molecules shown in panel **B**. A molecule is enriched inside hSSB1 droplets if the mentioned signal ratio is significantly higher than unity. Means ± SEM are shown together with individual data points of 10 images from *n* = 3 independent measurements.

Furthermore, hSSB1 was shown to assist the repair of ROS-induced DNA damage by recruiting hOGG1 to chromatin (*8*). Our fluorescence microscopy experiments showed that the mentioned proteins, and also hSSB2, were readily enriched inside LLPS droplets formed by hSSB1 (**Fig. 5B-C**). The SOSS component INIP, which only forms interactions with INTS3 but not with hSSB1 (*20*), showed enrichment in hSSB1 droplets only in the presence of INTS3 (**Fig. 5B-C**). In lack of specific interactions with hSSB1, neither the protein EGFP nor the small molecule fluorescein become enriched in condensates (**Fig. 5B-C**), indicating strong regulation regarding the contents of hSSB1 LLPS droplets. To rule out that other proteins are enriched inside hSSB1 droplets solely due to disulfide crosslinking resulting from the applied oxidizing conditions, we tested whether hSSB2, BLM, and INTS3 can be enriched in the droplets formed by the C41S-C81S-C99S variant hSSB1 protein (**Fig. S11A**). We observed similar coacervation to that seen with WT hSSB1. Furthermore, we tested whether the presence of ssRNA instead of ssDNA in hSSB1 droplets influences the enrichment of interaction partners, thus contributing to selectivity, but we found no such difference (**Fig. S11B**). These data show that hSSB1 condensates exert molecular filter functions by selectively recruiting and retaining interaction partners, similar to what we observed for the EcSSB protein (*23*).

### Oxidative stress induces hSSB1 organization into cytoplasmic foci that colocalize with stress granules but not with P-bodies

After our *in vitro* observation of redox-sensitive LLPS by hSSB1 in the presence of either ssDNA or ssRNA, we sought to explore the possibility of intracellular condensation of endogenous hSSB1 in HeLa cells under oxidative stress. Albeit hSSB1 has previously been implicated in nuclear functions (*5*, *8*, *13*, *15*, *17*, *21*, *22*), we detected its presence throughout the cell, both in the nucleus and cytoplasm, with the exception of the nucleolus (**Fig. 6A**). In untreated cells, the majority of hSSB1 is found in the nucleus, but a discernible fraction is located in the cytoplasm (**Fig. 6A**). Under these conditions, the cytoplasmic hSSB1 fraction showed disperse distribution with no apparent local enrichment or condensation (*8*).

**Fig. 6.**
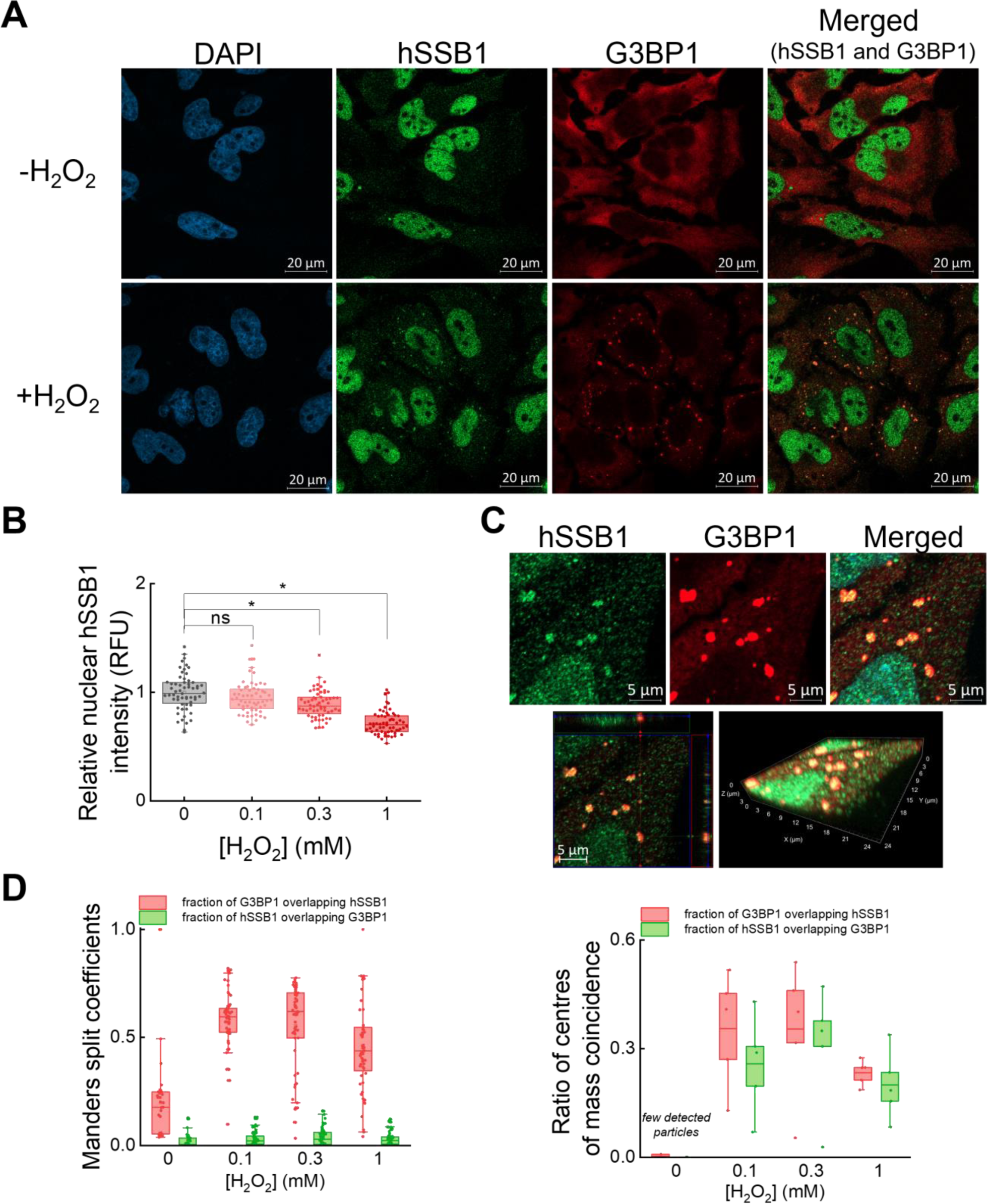
hSSB1 is organized into cytoplasmic foci upon oxidative stress, colocalizing with stress granules. (**A**) Representative confocal microscopy images (*n* = 3 independent experiments) of immunostained, untreated or H_2_O_2_-treated (300 µM, 2 h) HeLa cells. Blue channel shows nuclear DAPI stain, green and red channels show endogenous hSSB1 and G3BP1 stress granule marker, respectively. Merge image was created from hSSB1 and G3BP1 channels; yellow cytoplasmic foci indicate enrichment of hSSB1 in G3BP1-positive SGs. (**B**) Relative nuclear hSSB1 intensity of immunostained HeLa cells decreases upon H_2_O_2_ treatment (Mann-Whitney-test; * indicates significance; ‘ns’, not significant; *p* < 0.05; RFU, relative fluorescence units). Dots indicate individual nuclear intensities. (**C**) Top row, magnified confocal images of a H_2_O_2_-treated HeLa cell (cf. panel **A**), with hSSB1 and G3BP1 showing colocalized enrichment in cytoplasmic stress granules. Bottom row, 3D reconstructed images of the same ROI from confocal Z-stack images (16 slices, 3.6 μm optical sectioning). Green and red channels show endogenous hSSB1 and G3BP1 stress granule marker, respectively; yellow color indicates colocalization. (**D**) Manders split coefficients (left) and object based colocalization (right) of detected cytoplasmic hSSB1 and G3BP1 condensates. Dots indicate individual data derived from an image.

Surprisingly, we detected a significant, dose-dependent decrease in nuclear hSSB1 fluorescence intensity upon H_2_O_2_ treatment in immunocytochemical experiments (**Fig. 6B**). Moreover, we observed that upon acute oxidative stress caused by H_2_O_2_ treatment, endogenous hSSB1 became organized in multiple dense cytoplasmic foci in the size range of < 1 µm (**Fig. 6A**). Our *in vitro* measurements confirmed the ability of the applied antibodies to enter and stain hSSB1 droplets, thus enabling immunostaining of hSSB1 under LLPS conditions (**Fig. S12**). Stress granules (SGs) have been identified as cytoplasmic, membraneless ribonucleoprotein granules that form upon rapid changes in intra- or extracellular conditions, including oxidative stress (*29*, *30*). SGs exhibit liquid-like properties and were implicated in fine-tuning protein expression to adapt to changed environments. Ras GTPase-activating protein-binding protein 1 (G3BP1) was shown to be a major factor for SG assembly (*31*); therefore, we used G3BP1 staining to visualize SGs and assess their colocalization with hSSB1 granules. Diffuse G3BP1 staining with no SG formation was observed in untreated cells (**Fig. 6A**). However, we were able to trigger SG formation upon acute H_2_O_2_-induced oxidative stress (**Fig. S13A-B**) and observed strong colocalization between cytoplasmic hSSB1 granules and SGs (**Fig. 6C-D**). Manders split coefficients for the fraction of G3BP1 overlapping with hSSB1 indicated robust colocalization. However, Manders coefficients for the fraction of hSSB1 overlapping with G3BP1 appeared to be lower. This resulted from different levels of enrichment of G3BP1 and hSSB1 in SGs compared to the rest of the cytoplasmic intensity. Nevertheless, SG-independent hSSB1 granulation was scarcely seen, also supported by the ratio of object based colocalization of droplets (**Fig. 6D**). Furthermore, similar to the tested genome maintenance proteins, recombinant G3BP1 was also able to enrich inside hSSB1 droplets *in vitro*, in the presence of either ssRNA or ssDNA (**Fig. 5B-C, Fig. S11B**). Based on these findings, we conclude that hSSB1 is enriched inside SGs and colocalizes with the G3BP1 SG marker.

Paquet et al. demonstrated that hSSB1 robustly oligomerizes in U2OS cells upon oxidative stress (*17*). Therefore, we examined the extent of cellular hSSB1 covalent oligomerization at H_2_O_2_ concentrations that led to foci formation in HeLa cells (**Fig. S13C- D**). Upon H_2_O_2_ treatment, we observed a modest but significant increase in the dimer/monomer ratio of hSSB1 in whole cell lysates, while no higher order oligomers were seen. The extent of covalent dimerization in these experiments was lower compared to either that observed in our *in vitro* oxidation experiments (**Figs. 4C-D, S8**) or that reported by Paquet et al. (*17*) where robust hSSB1 covalent oligomerization was seen upon H_2_O_2_ treatment in U2OS cells. Notably, however, the extent of covalent dimerization aligns well with our observation that a discernible but small fraction of the total cellular hSSB1 pool localizes to SGs in HeLa cells (**Fig. 6A**). As we demonstrate that covalent oligomerization *per se* is neither a prerequisite for LLPS, nor is it alone sufficient for redox-dependent condensation (**Fig. 4**), we propose that in the cellular context, redox-dependent hSSB1 condensation may either be triggered by a small oxidized, but not necessary covalently linked fraction of hSSB1 oligomers, or be controlled by additional factors to be identified in further studies.

We also assessed hSSB1 subcellular patterns by transiently overexpressing EGFP- fused hSSB1 (hSSB1-EGFP) in HeLa cells. Interestingly, cells overexpressing hSSB1-EGFP showed granulation of both hSSB1-EGFP and G3BP1 even in the absence of oxidative stress, which was not seen in non-transfected cells (**Fig. S14A, cf. Fig. 6A**). Therefore, we investigated whether overexpression of hSSB1-EGFP, or that of EGFP alone, can influence SG formation. Assessed as a function of H_2_O_2_ concentration, we found the fraction of SG- containing cells to be similar in both EGFP- and hSSB1-EGFP expressing cells, with hSSB1- EGFP foci colocalizing with G3BP1 (**Fig. S14B-C**). While indicating that the transfection procedure and/or EGFP expression *per se* induce SG formation, in line with recent results suggesting that EGFP expression causes oxidative stress (*32*), these experiments corroborated the inclusion of hSSB1 in stress granules.

Processing-bodies (P-bodies) are membraneless ribonucleoprotein compartments associated with RNA degradation processes and share a number of components with SGs (*33*, *34*). However, while SGs are absent under normal conditions, P-bodies are constitutively present and observable at low levels. We used co-staining of P-body marker SK1-Hedls (*35*) and hSSB1 to investigate whether hSSB1 droplets colocalize with P-bodies under normal and oxidative stress conditions (**Fig. S15**). We observed no colocalization between the condensates, therefore we concluded that hSSB1 is not included in P-bodies.

### Stress granule associated hSSB1 condensation is accompanied by a decrease in nuclear hSSB1 levels in various cell lines under oxidative stress

It has been reported that hSSB1 relocalizes to chromatin after acute oxidative stress (*8*). Our experiments aimed to observe cytoplasmic granulation showed no nuclear enrichment, but a significant decrease in nuclear hSSB1 signal after 2 h of H_2_O_2_ treatment (**Fig. 6A-B**). Since nuclear accumulation was previously observed after 0.5 h of H_2_O_2_ treatment (*8*), we investigated the time dependence of cytoplasmic hSSB1 granulation and nuclear enrichment (**Fig. 7A**). Intriguingly, we found that 0.5 h of H_2_O_2_ treatment indeed causes a significant increase in nuclear hSSB1 intensity levels, but after 2 hours the nuclear fraction significantly decreased compared to control (**Fig. 7B**). Conversely, stress granule formation accompanied by cytoplasmic hSSB1 condensation was observed at low levels even after 0.5 h, and became robust after 2 h (**Fig. 7C**). Manders colocalization between G3BP1 and hSSB1 also appeared higher after 2 h of H_2_O_2_ treatment (**Fig. 7D**). Since the mentioned experiments were carried out in serum-free media, we tested whether acute serum deprivation causes similar phenotypes (**Fig. S16A**). We observed no cytoplasmic granulation, but a time-dependent decrease in nuclear hSSB1 fraction, which was restored after 4 h of serum deprivation (**Fig. S16B**). Changes in nuclear hSSB1 fractions during oxidative stress were thus compared to appropriate serum deprived controls throughout the article. Based on our results, we conclude that oxidative stress causes rapid accumulation of hSSB1 in nuclei, which is then depleted concomitantly with cytoplasmic accumulation. Other nuclear phenotypic changes, such as condensation similar to that seen in the cytoplasm, were not observed upon oxidative stress or serum deprivation. Small dense hSSB1 foci were discernible in nuclei (*e.g.* **Fig. 7A**, green channel), but these were permanently present and did not respond to any applied treatment reported in the present work.

**Fig. 7.**
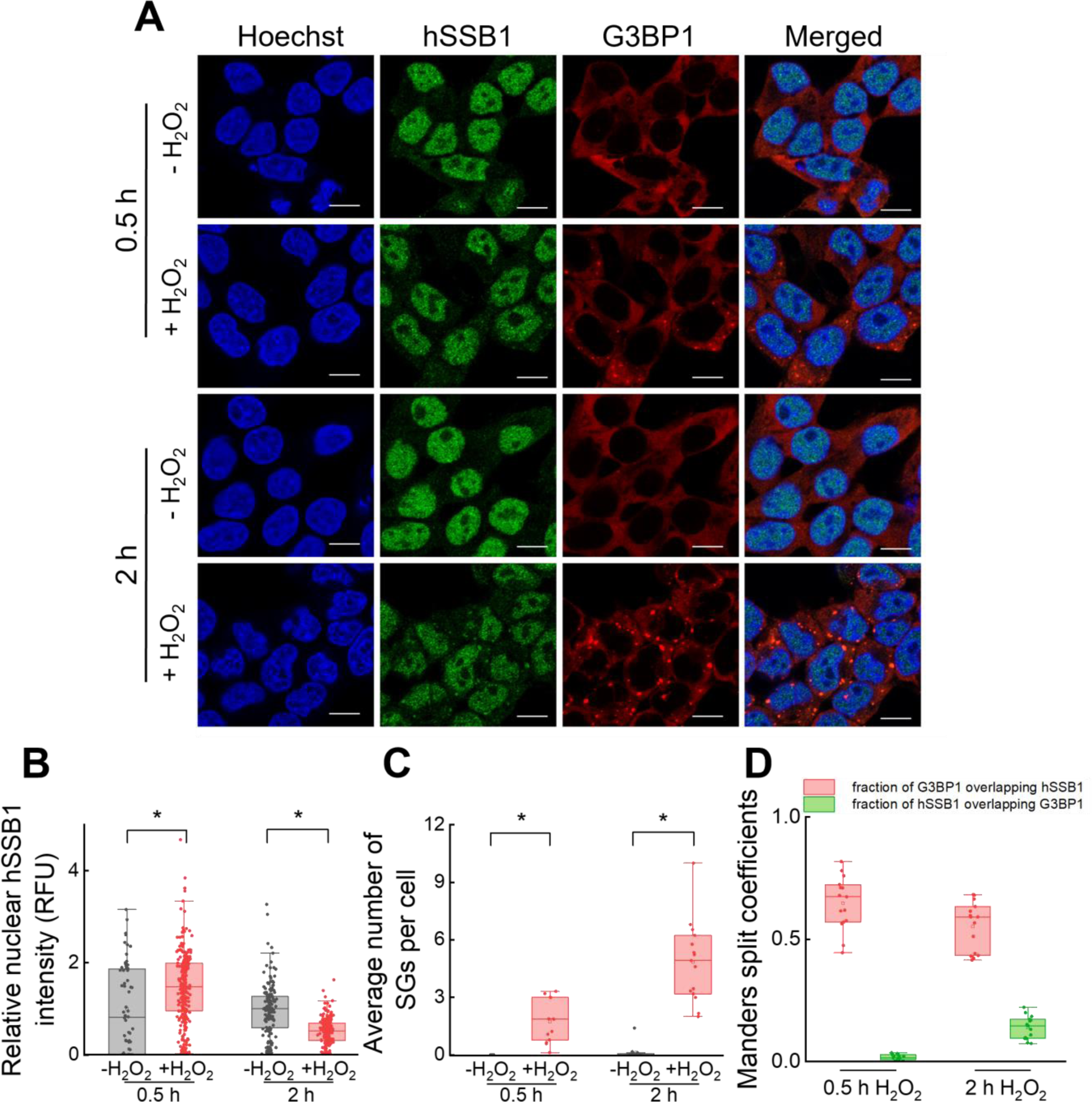
Oxidative stress triggers rapid nuclear accumulation of hSSB1, followed by cytoplasmic accumulation and stress granule associated hSSB1 condensation. (**A**) Representative confocal microscopic images (*n* = 3 independent experiments) of immunostained HeLa cells treated with 1 mM H_2_O_2_ for 0.5 h or 2 h. Blue channel shows nuclear Hoechst stain, green and red channels show endogenous hSSB1 and G3BP1 stress granule marker, respectively. Merged image was created from all three channels; yellow cytoplasmic foci indicate enrichment of hSSB1 in G3BP1-positive SGs. Scale bar: 10 μm. (**B**) Treatment with 1 mM H_2_O_2_ resulted in rapid (0.5 h) increase in nuclear hSSB1 signal (RFU, relative fluorescence units), followed by a decrease in nuclear hSSB1 intensity (2 h). As experiments were carried out in serum-free media, changes in nuclear hSSB1 intensities are normalized and shown next to appropriate serum deprivation controls (-H_2_O_2_) (see **Fig. S16**). Dots indicate individual nuclear intensities (Mann-Whitney-test, * indicates significant difference, *p* < 0.05). (**C**) Treatment with 1 mM H_2_O_2_ resulted in monotonously increasing number of SGs per cell over time. Dots represent average SG numbers derived from individual images (Mann-Whitney-test, * indicates significant difference, *p* < 0.05). (**D**) Manders split coefficients show colocalization of hSSB1 and G3BP1 condensates upon H_2_O_2_ treatment. Dots represent Manders oefficients derived from individual images.

Next, we aimed to investigate whether oxidative stress induced, SG-associated hSSB1 granulation can also be observed in other cell lines. Besides HeLa, we treated HEK293T and HFF-1 cells with H_2_O_2_ (**Fig. 8A**). In each case we observed a significant decrease in nuclear hSSB1 intensity after 2 h of treatment (**Fig. 8B**), accompanied by a significant increase in the number of SGs per cell (**Fig. 8C**). Interestingly, HeLa and HEK293T showed formation of a small number of relatively large SGs, while HFF-1 displayed a large number of much smaller SGs. In each cell line, SGs showed colocalization with hSSB1 condensates (**Fig. 8D**). Based on these results, we conclude that oxidative stress causes transient nuclear hSSB1 depletion and gradual, SG-associated hSSB1 condensation in both cancerous and non-cancerous cell lines, suggesting that redox-dependent hSSB1 LLPS can occur in cells with different metabolic profiles.

**Fig. 8.**
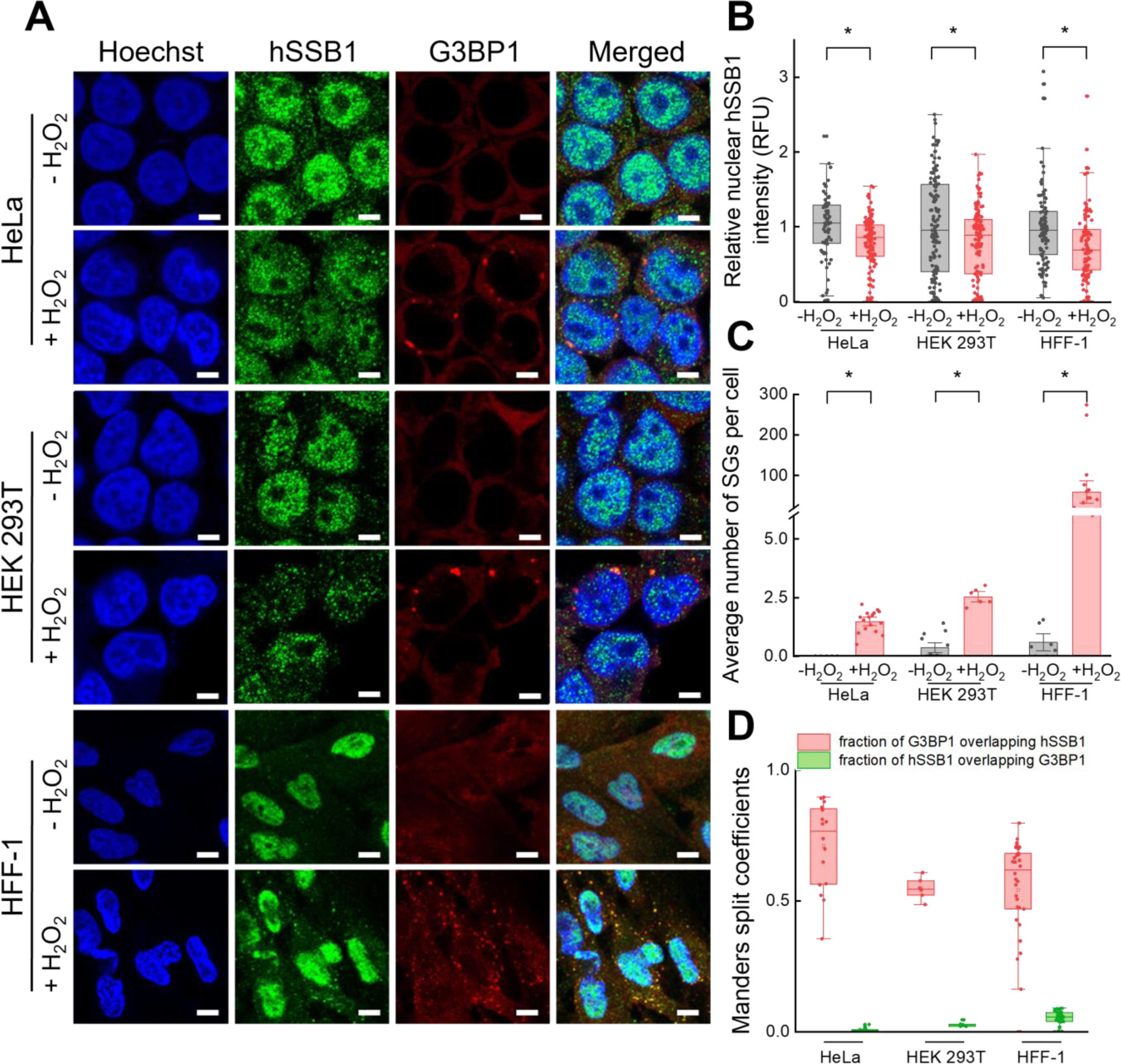
Stress granule associated hSSB1 condensation is accompanied by nuclear hSSB1 intensity decrease in various cell lines under oxidative stress (**A**) Representative confocal microscopic images (*n* = 3 independent experiments) of immunostained HeLa, HEK293T and HFF-1 cells treated with H_2_O_2_ (1 mM, 2 h). Blue channel shows nuclear Hoechst stain, green and red channels show endogenous hSSB1 and G3BP1 stress granule marker, respectively. Merge image was created from all three channels; yellow cytoplasmic foci indicate enrichment of hSSB1 in G3BP1-positive SGs. Scale bar: 5 μm. (**B**) H_2_O_2_ treatment (1 mM, 2 h) causes a decrease in nuclear hSSB1 intensity in each cell line. As experiments were carried out in serum-free media, changes in nuclear hSSB1 intensities (RFU, relative fluorescence units) were normalized and shown next to appropriate serum deprivation controls (-H_2_O_2_). Dots indicate individual nuclear intensities (Mann- Whitney-test, * indicates significant difference, *p* < 0.05). (**C**) H_2_O_2_ treatment (1 mM, 2 h) induces robust SG formation in each cell line. Dots represent average SG numbers derived from individual images (Mann-Whitney-test, * indicates significant difference, *p* < 0.05). (**D**) Manders split coefficients show colocalization of hSSB1 and G3BP1 condensates upon H_2_O_2_ treatment (1 mM, 2 h) in each cell line. Dots represent Manders coefficients derived from individual images.

### hSSB1 is enriched in stress granules in response to various forms of cellular stress

Subsequently, we investigated the effect of a variety of cellular stressors on hSSB1 condensation (**Fig. 9A**). Besides H_2_O_2_, we treated HeLa cells with potassium bromate (KBrO_3_) and sodium arsenite (NaAsO_2_), which are often used to induce direct oxidative stress (*17*, *31*). Menadione was shown to generate ROS through redox cycling while disrupting mitochondrial membrane potential, triggering cytochrome c redistribution to the cytosol and inducing cell death, thus inducing indirect, internal oxidative stress (*36*). DTT was used to induce endoplasmic reticulum stress by inhibiting protein folding through the reduction of disulfide bridges (*37*). DTT is also responsible for ROS production by influencing cell signaling and the glutathione system (*37*). Treatment with NaAsO_2_, menadione sodium bisulfite, and DTT, similarly to H_2_O_2_, resulted in the decrease of nuclear hSSB1 signal compared to appropriate serum deprivation controls according to treatment time (**Fig. 9B**), accompanied by SG associated hSSB1 granulation (**Fig. 9C-D**). Interestingly, no phenotypic changes were observed in response to KBrO_3_. These stressors exert their effect mainly in the cytoplasm, thus we wished to investigate an agent that is genotoxic and disrupts DNA metabolism. Etoposide inhibits DNA topoisomerase II, thus leading to the accumulation of DNA breaks, at which hSSB1 was shown to localize (*13*, *15*, *38*). Unexpectedly, no nuclear hSSB1 accumulation or condensation was observed in response to etoposide treatment.

**Fig. 9.**
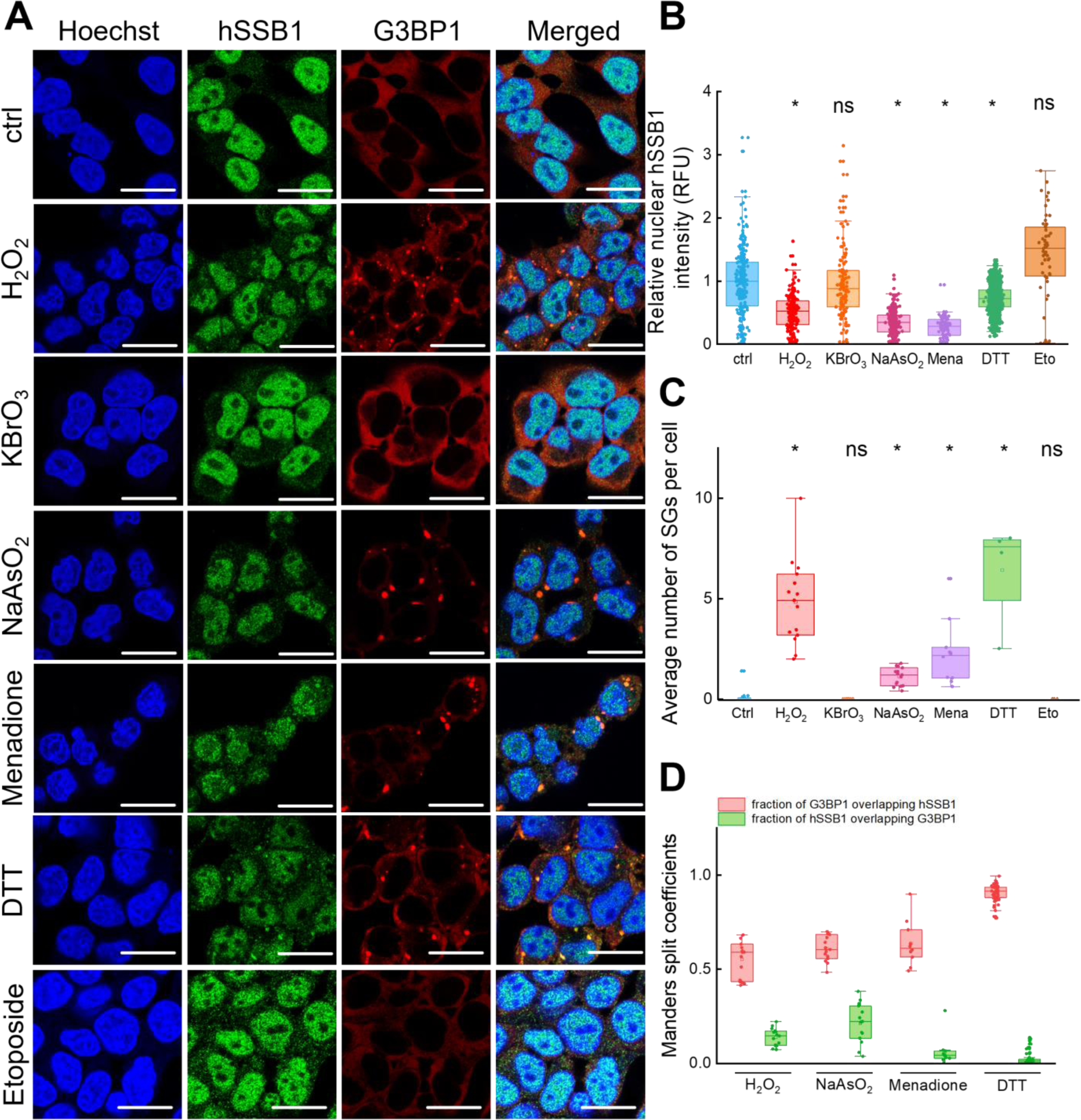
hSSB1 is enriched in stress granules in response to various cellular stressors. (**A**) Representative confocal microscopic images (*n* = 3 independent experiments) of immunostained HeLa cells treated with various stress agents (H_2_O_2_: 1 mM, 2 h; KBrO_3_: 30 mM, 2 h; NaAsO_2_: 0.5 mM, 2 h; Menadione sodium bisulfite: 100 μM, 4 h; DTT: 1 mM, 2 h; Etoposide: 100 μM, 2 h). Blue channel shows nuclear Hoechst stain; green and red channels show endogenous hSSB1 and G3BP1 stress granule marker, respectively. Merged image was created from all three channels; yellow cytoplasmic foci indicate enrichment of hSSB1 in G3BP1-positive SGs. Scale bar: 20 μm. (**B**) Besides H_2_O_2_, NaAsO_2_, menadione sodium bisulfite (mena), and DTT triggered a decrease in nuclear hSSB1 intensity in HeLa cells, while KBrO_3_ and etoposide (eto) did not. As experiments were carried out in serum-free media, changes in nuclear hSSB1 intensities (RFU, relative fluorescence units) were normalized to appropriate serum deprivation controls. Dots indicate individual nuclear intensities (Kruskal-Wallis test with Dunn’s post-hoc test, * indicates significant difference, ‘ns’ not significant, *p* < 0.05). (**C**) Besides H_2_O_2_, NaAsO_2_, menadione sodium bisulfite, and DTT induced robust SG formation, while KBrO_3_ and etoposide did not. Changes in the average number of SGs per cell are shown. Dots represent average SG numbers derived from individual images (Kruskal-Wallis test with Dunn’s post-hoc test, * indicates significant difference, *p* < 0.05). (**D**) Manders split coefficients show colocalization of hSSB1 and G3BP1 condensates upon treatment with H_2_O_2_, NaAsO_2_, menadione sodium bisulfite, and DTT. Dots represent Manders coefficients derived from individual images.

Nevertheless, taken together our data show that various stress agents affecting redox conditions, protein folding, and ROS signaling induce cytoplasmic, SG associated granulation of hSSB1.

### hSSB1 silencing enhances stress granule formation upon oxidative stress

After establishing that hSSB1 shows SG-associated cytoplasmic condensation during cellular stress response, we set out to examine the effect of hSSB1 silencing on SG formation. hSSB1 silencing was achieved by lipofection of HeLa cells with a pool of siRNAs (see Methods).

Appropriate vehicle and non-targeting RNA controls were used. hSSB1 silencing was verified by Western blot, and silencing efficiency was determined to be ∼70 % (extent of reduction in hSSB1 protein content) based on GAPDH loading control (**Fig. 10A**). Confocal images of immunostained HeLa cells showed robust decrease in nuclear hSSB1 intensity upon silencing (**Fig. 10B**). Upon oxidative stress, SGs were visible in the cytoplasm even when hSSB1 was silenced. Moreover, the residual hSSB1 fraction still showed discernible enrichment inside G3BP1-stained SGs (**Fig. 10B**). In hSSB1-silenced cells, nuclear hSSB1 intensity was around 3-fold lower compared to control cells, but the oxidative stress-induced nuclear depletion remained unchanged (**Fig. 10C**). SG formation in case of non-transfected control cells and non-target controls upon H_2_O_2_ -treatment was similar to that seen previously. Surprisingly, hSSB1 silencing resulted in a significantly higher fraction of SG+ cells after 0.3 mM H_2_O_2_ treatment, compared to non-silenced controls (**Fig. 10D**). This result suggests that hSSB1 may play a stress dose dependent regulatory role in SG formation.

**Fig. 10.**
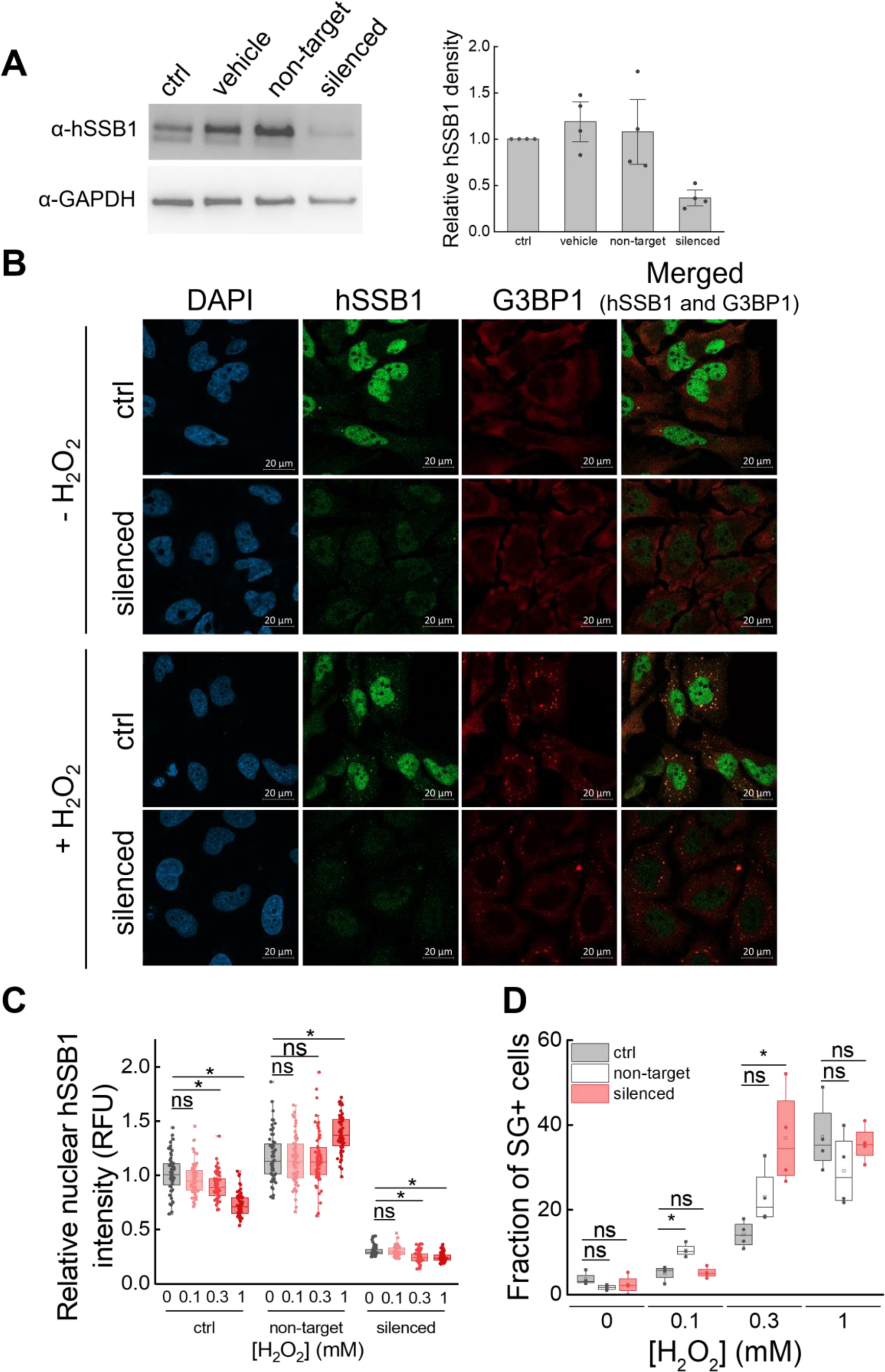
Effect of hSSB1 silencing on stress granule formation. (**A**) Representative Western blots (left) analyzed by pixel densitometry (right; dots indicate individual experiments) show successful silencing of hSSB1, using GAPDH loading control. ’Ctrl’ indicates no transfection, ’vehicle’ represents immunoblots from cells transfected with empty liposomes, while ’non-target’ samples were exposed to liposomes containing non- targeting siRNA (see Methods). Error bars represent SEM. Means are shown normalized to untransfected control for each dataset. (**B**) Representative confocal microscopic images (*n* = 3 independent experiments) of immunostained HeLa cells silenced for hSSB1, shown next to unsilenced controls (ctrl). Cells were treated with H_2_O_2_ (0.3 mM, 2 h) as indicated. Blue channel shows nuclear DAPI stain, green and red channels show endogenous hSSB1 and G3BP1 stress granule marker, respectively. Merged image was created from hSSB1 and G3BP1 channels; yellow cytoplasmic foci indicate enrichment of hSSB1 in G3BP1-positive SGs. (**C**) H_2_O_2_ causes a decrease in nuclear hSSB1 intensity (cf. **Fig. 7B**). Silencing greatly lowers nuclear hSSB1 intensity, while not affecting stress-induced change in distribution. Nuclear hSSB1 intensities (RFU, relative fluorescence units) were normalized to untreated, untransfected controls. Dots indicate individual nuclear intensities (Mann-Whitney-test, * indicates significant difference, ‘ns’ not significant, *p* < 0.05). (**D**) Immunocytochemistry- based cell typization (fraction of SG+ cells) showing the effects of H_2_O_2_ treatment and hSSB1 knockdown. hSSB1 silencing renders cells more susceptible to oxidative stress-induced SG formation at 0.3 mM (2 h) H_2_O_2_ treatment. Dots indicate data collected from subimages (see Methods) (two-sample T-tests, * indicates significant difference, ‘ns’ not significant, *p* < 0.01).

## DISCUSSION

In this work we show that hSSB1 forms phase-separated liquid condensates under physiologically relevant macromolecular and ionic conditions *in vitro* and shows cytoplasmic, SG-associated granulation upon various forms of cellular stress in both cancerous and non-cancerous cell lines. hSSB1 coacervates with nucleic acids, both DNA and RNA, in a stoichiometry-dependent fashion, with its condensation being selectively initiated by oxidative conditions (**Figs. 2-3, S3-6**). hSSB1 condensation is effective even at low protein concentrations, without the need for molecular crowders (**Fig. 3A-B**) and is driven by hSSB1’s intrinsically disordered region (IDR) (**Figs. 4A, S7**), in line with PDB-deposited crystal structures obtained with a truncated hSSB1 construct supporting the propensity for IDR-IDR intermolecular interactions (PDB codes 5D8E and 5D8F, no accompanying publication) (**Fig. 11A**).

**Fig. 11.**
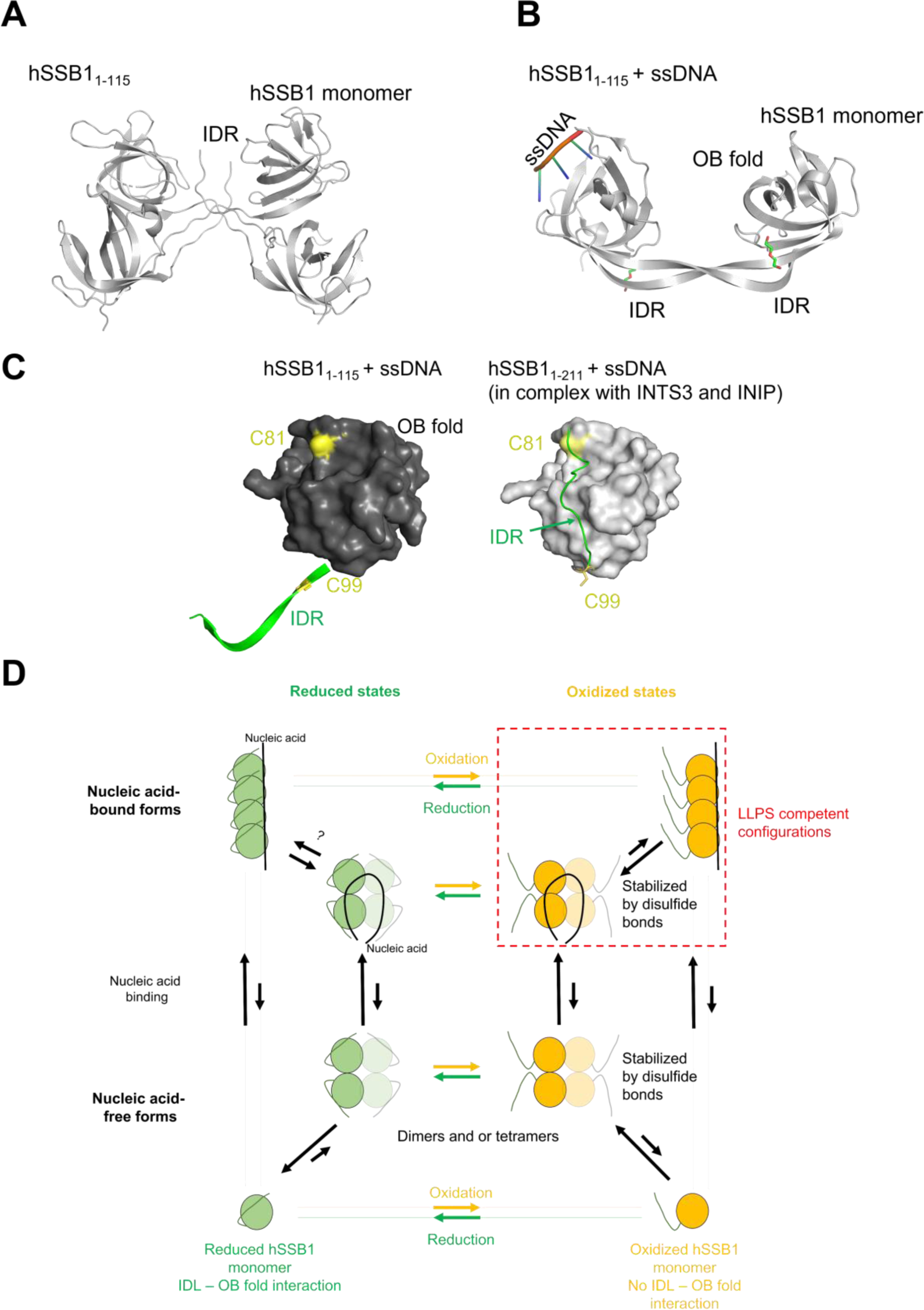
Proposed model for the role of hSSB1 IDR tail flexibility in redox-dependent LLPS regulation. (**A-B**) Crystal structures of a truncated hSSB1 construct (amino acids 1- 115) lacking the majority of the IDR, in the absence (**A**) and presence of ssDNA (**B**) (PDB codes 5D8E and 5D8F, respectively). In both conditions, interactions can be observed between the N-terminal IDR segment contained in the constructs. (**C**) Comparison of the structure of ssDNA-bound truncated hSSB1 (dark grey, PDB code 5D8E, cf. panel **B**) and that of full-length hSSB1 in the ssDNA-bound SOSS1 complex (light grey, PDB code 4OWW; INTS3 and INIP structures were removed for clarity) (*19*). The N-terminal part of the IDR is not resolved in the latter structure. For both structures, the OB domain is shown as a surface model and the IDR tails are represented as green cartoons. Note the large difference between IDR conformations. (**D**) Proposed model for regulation of hSSB1 LLPS. In the reduced form and in the absence of nucleic acids, hSSB1 is in a dynamic equilibrium between monomeric and oligomeric states, with the monomeric form being favored (*6*, *17*). Binding to single-stranded nucleic acids may either stabilize/facilitate protein oligomerization, or hSSB1 monomers can bind independently to the nucleic acid lattice in a sequential order without formation of oligomers. Under reducing conditions, the C-terminal part of the hSSB1 IDR can interact with the OB fold (cf. panel **C**, right side) both in the absence and presence of nucleic acids. Upon oxidation, the IDR is released from the OB fold and participates in intermolecular IDR-IDR interactions (cf. panel **C**, left side). Covalent hSSB1 oligomers form via disulfide bridges involving C41 and C99; however, covalent oligomerization is not required for LLPS *per se* (cf. **Fig. 4D-E**). LLPS is triggered by multivalency brought about by nucleic acid binding (independent of hSSB1 oligomerization state) and oxidation-dependent, conformationally regulated IDR-IDR interactions.

Importantly, the dependence of hSSB1 LLPS on oxidative conditions (**Fig. 2, Fig. S3**) strongly suggests a cellular oxidative stress sensing/response role for the protein. Effects caused by ROS are thought to be exerted locally (*39*, *40*), where hSSB1 droplets, as first responders, could rapidly form on exposed single-stranded nucleic acid segments.

We also show that other genome maintenance proteins interacting with hSSB1 can readily and selectively be enriched in hSSB1 droplets (**Figs. 5, S11**). hSSB1 condensates can thus recruit required factors to the site of action, and also act as a molecular filter to restrict the localization of other proteins. One such example may be BER where hSSB1 was shown to assist the repair of ROS-induced DNA damage by recruiting hOGG1 to chromatin (*8*).

However, considering that we observed hSSB1-driven LLPS also in conjunction with ssRNA (**Fig. 2A**), the roles of hSSB1 LLPS may not be limited to genome repair, as it may also be effective in RNA metabolic processes both in the nucleus and in the cytoplasm (see below).

While intermolecular IDR-IDR interactions appear to be a major driving force for LLPS (**Fig. 4A**), binding of single-stranded nucleic acids is also crucial for coacervation (**Fig. 2A, 3C**). hSSB1 was shown to exist dominantly as a monomer in the absence of nucleic acids (*6*, *17*). However, as LLPS processes generally require multivalent interactions, we propose that single-stranded nucleic acids can either act as a scaffold to which multiple hSSB1 monomers can bind sequentially and independently in a beads on a string-like fashion or as a stabilizer for oligomerization. In either case, the multivalency of SSB-SSB interactions is enhanced in addition to the increased local concentration of SSB IDR regions. The bell-shaped ssDNA concentration dependence of hSSB1 LLPS (**Figs. 3C-D, S6**) can be explained based on the beads on a string-like model, regardless of the oligomerization state of the protein. When the concentration of ssDNA molecules exceeds the SSB binding stoichiometry, the probability of multiple SSB molecules binding to the same ssDNA molecule decreases, leading to a lower LLPS propensity. However, our findings that despite significant hSSB1 oligomerization upon oxidation in the absence of nucleic acids (**Fig. 4C-D**), LLPS is only observed upon nucleic acid binding, indicate additional unexplored roles for nucleic acid binding, potentially distinct from the scaffolding function.

The role of protein oxidation in LLPS regulation is especially intriguing. According to an emerging concept, cysteine residues within proteins may act as regulatory switches in redox signaling (*41*, *42*). In this work we comprehensively assessed the roles of each of hSSB1’s cysteine residues in the interplay between covalent oligomerization and nucleoprotein condensation (**Figs. 4C-E, S8-9**). On one hand, we show that covalent oligomerization of hSSB1, mediated by residues C41 and C99, is not a prerequisite for forming LLPS condensates (**Fig. 4D**). On the other hand, the capability for covalent oligomerization alone is not sufficient for redox-dependent LLPS, for which all three hSSB1 cysteines are required (**Fig. 4E**). Taken together, our observations point toward a delicate autoregulatory mechanism for LLPS, which apparently involves oxidation-dependent hSSB1 structural changes. While the structure of oxidized hSSB1 is yet unknown, existing crystal structures also support a regulation model as follows. In both of nucleic acid-free and ssDNA-bound structure of a truncated hSSB1 construct lacking the majority of the IDR (PDB codes 5D8E and 5D8F), extensive interactions between the remaining C-terminal parts of the tail are observed (**Fig. 11A-B**). In contrast, in the SOSS1 (INTS3, INIP, SSB1) complex, the same region of full-length hSSB1 folds back onto the OB domain both in the presence and absence of ssDNA (*19*). Moreover, the N-terminal part of the IDR is not resolved, probably because it is disordered (**Fig. 11C**). The large difference in protein structure, i.e., the position of the IDR tail, in the above-mentioned structures highlights the mobility of the tail and its capability to either bind to the OB fold intramolecularly or to interact with other hSSB1 tails (**Fig. 11C**).

Importantly, in the OB domain-bound form of the IDR, its C-terminal part lies in a valley on the OB domain’s surface, which includes cysteine 81 and a hydrophobic pocket adjacent to C99 (**Fig. 11C**). Based on these observations, we can envision a scenario in which the oxidation of these residues (C81, C99), and also that of cysteine 41, could potentially alter the structure of hSSB1 and significantly weaken the interaction between the IDR and the OB fold. This effect may facilitate intermolecular IDR-IDR interactions that are essential for LLPS. The observed IDR-IDR and IDR-OB interactions are apparently mutually exclusive as they involve the same residues and likely compete with each other based on the structural data.

Our results suggest that all three cysteines in their reduced form contribute to the inhibition of interprotein interactions essential for LLPS, regardless of the presence of nucleic acids, possibly by facilitating IDR-OB fold interactions and thus inhibiting IDR-IDR interactions. Single or combinatorial mutations of any of the cysteines eliminate the inhibitory effect and thus abolish redox sensitivity of hSSB1 LLPS (**Fig. 4E**). In addition, cysteines 99 and 41 are involved in the formation of covalently bound oligomers. While covalent oligomers are not required for hSSB1 LLPS, they were shown to bind ssDNA with enhanced affinity compared to the monomeric form, indicating a role in regulating interactions within and stability of the nucleoprotein complex (*43*). In addition, disulfide bridges may stabilize the IDR in an OB-unbound form in hSSB1 oligomers. Based on the above findings, we propose a model for a regulatory mechanism that ensures that robust LLPS is only triggered when the hSSB1 protein is oxidized and bound to nucleic acids (**Fig. 11D**). Although redox-dependent regulatory modifications (*42*) are being identified in an increasing number of proteins (*44*), evidence is still scarce for redox-sensitive amino acid modifications governing LLPS propensity. Examples include disulfide formation-dependent LLPS by TMF (terminating flower) transcription factor, governing plant tissue development in the apical meristem (*45*), and LLPS by yeast Ataxin-2 mediated by reversible oxidation of methionine (*46*). Further exploration of redox-dependent molecular changes in hSSB1 nucleoprotein coacervates will likely yield key insights into mechanisms of oxidative stress response.

As hSSB1 and hSSB2 show high structural resemblance (**Fig. 1A**), recently we also investigated the LLPS propensity of hSSB2 *in vitro* (*47*). Similar to hSSB1, hSSB2 also undergoes LLPS upon ssDNA binding in physiologically relevant ionic conditions at low protein concentrations, mediated by the IDR region. hSSB2 LLPS also shows a bell-shaped dependence on nucleic acid concentration. However, ssRNA only moderately enhances hSSB2 condensation compared to ssDNA, although hSSB2 binds to ssDNA and ssRNA with similar affinities. Interaction partners of hSSB2 become selectively enriched inside hSSB2 condensates. However, *bona fide* hSSB2 condensation was only seen in reducing conditions, while oxidation promoted the formation of branched, solid-like nucleoprotein particles. All cysteine residues are conserved between hSSB1 and hSSB2 (C45, C85, C103 in hSSB2); thus, the observed differences between hSSB1 and hSSB2 condensation indicate additional, yet unknown structural mechanisms of redox-sensitive LLPS regulation besides cysteine oxidation.

In the present study we found that a discernible fraction of the hSSB1 protein is present in the cytoplasm, which is organized into distinct granules upon acute oxidative stress (**Fig. 6A-B**), although this far only nuclear functions have been described for hSSB1.

Cytoplasmic hSSB1 foci colocalize with stress granules (**Fig. 6C-D**) but not with P-bodies (**Fig. S15**). Interestingly, hSSB1 has not been implicated to be me a member of the SG proteome by previous approaches aiming to identify SG constituents (*48*).

The previously reported increased H_2_O_2_ sensitivity of cells upon hSSB1 silencing was interpreted as a result of the defect in the repair of 8-oxoguanine DNA lesions through BER (*8*). However, our current findings suggest that hSSB1 plays additional roles in processes distinct from genome repair. We find that hSSB1 forms coacervates with ssRNA as effectively as with ssDNA (**Fig. 2A**). Thus, ssDNA regions appearing upon DNA damage can act as a scaffold to initiate hSSB1-driven LLPS in the nucleus, while in the cytoplasm ssRNA regions can facilitate droplet formation and incorporation of hSSB1 condensates into SGs. hSSB1 knockdown resulted in an elevation in the fraction of cells displaying SGs upon 0.3 mM H_2_O_2_ treatment (**Fig. 10D**), indicating a negative influence of hSSB1 on SG formation under these conditions. Interestingly, hSSB1 is not the only genome maintenance protein that has been shown to enter SGs. BLM helicase and hOGG1, interaction partners of hSSB1 (*8*, *15*), which we demonstrate to be enriched in hSSB1 granules *in vitro* (**Fig. 5**), were detected in SGs formed upon various forms of stress (*49*, *50*). BLM was found to inhibit SG formation *via* unwinding cytoplasmic RNA G-quadruplexes associated with cellular stress response. BLM knockdown resulted in increased SG formation, similarly to hSSB1 knockdown in our study. Considering that hSBB1 is needed for stability and recruitment of BLM at DNA double strand breaks and stalled replication forks (*15*), one can envision a similar scenario whereby hSSB1 could assist the recruitment of BLM to RNA quadruplexes inside SGs, further expanding the physiological implications of our results. Similarly, hSSB1 may facilitate the recruitment of hOGG1 to SGs.

We found that oxidative stress induced rapid nuclear accumulation of hSSB1, followed by cytoplasmic accumulation (**Fig. 7B**). Besides this effect, we did not detect redox-regulated nuclear hSSB1 condensation. As hSSB1 oligomerization is required for efficient hOGG1-mediated BER (*8*) and we demonstrated that hOGG1 is readily enriched inside hSSB1 condensates (**Fig. 5**) redox-dependent LLPS may play a role in the repair of oxidative DNA lesions. Recent evidence also suggests that hSSB1’s LLPS propensity may be modulated by interaction partners. INTS3 is a major nuclear interacting partner of hSSB1 in the SOSS1 complex (*21*, *51*), and it also contains a C-terminal IDR (*52*). The ternary SOSS1 complex was recently demonstrated to undergo LLPS, with condensates being localized to laser induced DSBs (*52*). The purified SOSS1 complex was able to undergo LLPS in the presence of molecular crowder, and droplet formation was enhanced upon addition of ssRNA or ssDNA. Truncation of the C-terminal IDR of INTS3 inhibited droplet formation by the SOSS1 complex *in vitro.* While the redox dependence of SOSS1 LLPS has not been tested, these findings indicate that the LLPS properties of hSSB1 are significantly modulated by its partners in the SOSS1 complex. LLPS by SOSS1 was also recently demonstrated to be involved in transcription regulation and the prevention of R-loop induced genome instability (*53*). Nuclear hSSB1 puncta were seen even in stress-free conditions, which resembled the small nuclear hSSB1 foci visible in our experiments both under stress-free and stress conditions (*e.g.* **Fig. 7A**, green channel). Taken together, these findings bear further implications for the physiological importance of hSSB1-driven nuclear LLPS transitions.

hSSB1 is not predicted to harbor any classical or non-classical nuclear localization signals (NLS) (*54*, *55*), and it has not been found to interact with importins. We propose that the small monomeric hSSB1 protein (23 kDa) may enter the nucleus *via* passive diffusion or attached to NLS-harboring interaction partners, e.g. INTS3 (*56*), while a cytoplasmic hSSB1 fraction associates with SGs, coupled to its LLPS being triggered upon cellular stress.

Besides hSSB1 and the SOSS1 complex, the well-known ssDNA-binding protein RPA has also recently been shown to undergo LLPS, and RPA droplets were shown to colocalize with telomeres (*57*). Similar to hSSB1 condensation, LLPS by RPA is enhanced by ssDNA. However, unlike that for hSSB1, ssRNA did not induce LLPS by RPA or become enriched inside condensates. Droplet formation by RPA is inhibited by phosphorylation, a regulatory mechanism also plausible for hSSB1 condensation, as hSSB1 is phosphorylated by multiple kinases (*5*, *52*, *58*). The spatial regulation and potential co-existence of hSSB1 and RPA condensates is yet to be elucidated.

The role of hSSB1 and its discovered LLPS propensity in oxidative stress response is particularly relevant in the context of cancer cells that generally experience chronic oxidative stress. It is plausible that hSSB1’s capability to form condensates in a redox-dependent manner supports cancer cell survival. Indeed, hSSB1 is upregulated in a large variety of cancers including gastric and colorectal adenocarcinoma, chronic lymphocytic leukemia, B-cell non-Hodgkin lymphoma, breast adenocarcinoma, and hepatocellular carcinoma (**Fig. S17**). In addition, hSSB1 was recently shown to be involved in DNA damage response and transcription regulation processes in prostate cancer (*59*). Considering that numerous anticancer agents act at least in part by generation of oxidative stress, targeting of hSSB1’s LLPS propensity appears as a promising tool to sensitize cancer cells to oxidative stress without compromising the LLPS-independent functions of the protein.

## MATERIALS AND METHODS

### General reaction conditions

Unless otherwise stated, *in vitro* measurements were performed at 25°C in LLPS buffer containing 25 mM Tris-HCl pH 7.4, 10 mM MgCl_2_, 50 mM KCl, supplemented with 0.003 % (980 µM) H_2_O_2_ or 1 mM DTT. LLPS buffer omitting MgCl_2_ and KCl (denoted as “no salt LLPS buffer”) was used where indicated. We note that the latter experiments contained 1.7 mM KCl and 3.4 mM MgCl_2_ originating from the storage buffer of hSSB1.

### Cloning, protein expression and purification

pET29a-hNABP2 (hSSB1), pET29a-hNABP1 (hSSB2), pET28a-C9ORF80, and pGEX-6P-1-INTS3-FL were gifts from Yuliang Wu (Addgene plasmids #128307, 128306, 128418, and 128415, respectively) (*6*, *51*). pET29a-hSSB1 contained a point mutation coding for an Y85C substitution, which was restored to WT tyrosine using QuikChange (Agilent) mutagenesis.

QuikChange results were verified by DNA sequencing. Expression and purification of hSSB1 and hSSB2 fused to a C-terminal histidine-tag were performed as described in references (*5*, *6*) with modifications as follows. In the case of hSSB1, Ni-NTA beads were equilibrated with buffer A (25 mM Tris-HCl pH 8.0, 500 mM NaCl, 10 % glycerol, 0.1 % TWEEN 20). Loaded column was washed with 10 column volumes of buffer B (25 mM Tris-HCl pH 8.0, 50 mM NaCl, 10 % glycerol, 0.1 % TWEEN 20) supplemented with 50 mM imidazole. Protein was eluted with buffer B containing 250 mM imidazole, and loaded on HiTrap Heparin column (GE Healthcare) equilibrated with HP1 buffer (25 mM Tris-HCl pH 8.0, 50 mM NaCl, 10 % glycerol). Elution was achieved with HP2 buffer (25 mM Tris-HCl pH 8.0, 1 M NaCl, 10 % glycerol). Purified protein samples were concentrated using Amicon Ultra 10K spin column (Sigma-Aldrich) and dialyzed against Storage buffer (25 mM Tris-HCl pH 8.0, 50 mM KCl, 100 mM MgCl_2_, 10 % glycerol, 1 mM DTT). Plasmids coding for hSSB1 variants were generated from the Y85C substitution-corrected pET29a-hSSB1 vector using the QuikChange (Agilent) mutagenesis kit, and hSSB1 variant proteins were purified as WT hSSB1. In case of hSSB1-IDR, a stop codon was introduced after the codon coding for aa 109. Mutagenesis was verified by DNA sequencing. In the case of hSSB2, only nickel affinity chromatography was applied.

INTS3 and INIP were purified as described (*51*) with modifications for INTS3 as follows. After sonication, GST-tagged INTS3 was loaded onto Glutathione Agarose (Pierce™) column equilibrated with buffer A (25 mM Tris-HCl pH 8.0, 500 mM NaCl, 1 mM EDTA, 1 mM DTT, 0.2 % Tween 20, 10 % glycerol). Loaded column was washed with 2 column volumes of buffer B (25 mM Tris-HCl pH 8.0, 150 mM NaCl, 1 mM DTT, 0.2 % Tween 20, 10 % glycerol). Precision protease (10 unit/ml) was introduced in buffer B on column and INTS3 was digested overnight. Tag-free protein was eluted with buffer B, then concentrated with Amicon Ultra 100K spin column (Sigma-Aldrich) and dialyzed against Storage buffer (25 mM Tris-HCl pH 8.0, 50 mM NaCl, 10 % glycerol, 1 mM DTT).

Plasmids, expression, and purification of EcSSB, human BLM and EGFP constructs were described in (*23*, *60*). Recombinant hOGG1 (ab98249) and G3BP1 (ab103304) were purchased from Abcam.

Purity of samples was checked by tris-glycine based gradient SDS-PAGE (Mini-Protean TGX 4-20 %, Bio-Rad) for all constructs. Bradford method was used for concentration measurement of proteins. Purified proteins were frozen in droplets and stored in liquid N_2_.

pEGFP-C1 plasmid, used for transfection of HeLa cells, was obtained from Clontech. The hSSB1 coding sequence was amplified by PCR and cloned upstream of EGFP to fuse hSSB1 to the N-terminus of EGFP. Constructs were verified by DNA sequencing.

### Fluorescent labeling of proteins

hSSB2 was labeled with 5-IAF (5-iodoacetamido-fluorescein, Thermo Fisher) on the intrinsic cysteines. hSSB2 Storage buffer (25 mM Tris-HCl pH 8.0, 50 mM NaCl, 100 mM MgCl_2_, 10% glycerol, 1 mM DTT) was exchanged to a storage buffer omitting DTT using a PD10 (GE Healthcare) gel filtration column. IAF was applied at 1.5-fold molar excess compared to hSSB2. The labeling reaction was performed for 3.5 h in argon atmosphere, at room temperature. The labeled protein was purified with a PD10 (GE Healthcare) gel filtration column pre-equilibrated with hSSB2 storage buffer and dialyzed against storage buffer to remove residual free dye.

INIP, hOGG1, and G3BP1 were labeled on their N-termini with AF488 (Alexa Fluor 488 carboxylic acid succinimidyl ester, Thermo Fisher). Storage buffer was exchanged to Labeling buffer (50 mM MES pH 6.5, 50 mM KCl, 10% glycerol) by dialysis. AF488 was introduced in 4-fold molar excess over protein. Labeling process was performed for 5 h at room temperature. Reaction was stopped by 50 mM Tris-HCl final concentration. For INIP, the labeled protein was purified using a PD10 (GE Healthcare) gel filtration column pre-equilibrated with INIP Storage buffer (25 mM Tris-HCl pH 8, 150 mM KCl, 10% glycerol). and was dialyzed against Storage buffer and repetitively filtered using an Amicon Ultra 10K spin column to remove residual free dye. Labeled hOGG1 and G3BP1 were repetitively filtered on Amicon Ultra 10K and 30K to exchange labeling buffer to storage buffers provided by manufacturers.

hSSB1 and INTS3 proteins were labeled with AF647 (Alexa Fluor 647 carboxylic acid succinimidyl ester, Thermo Fisher). Proteins were dialyzed against Labeling buffer (50 mM MES pH 6.5, 50 mM KCl, 100 mM MgCl_2_, 5 mM DTT). AF647 was applied at 1.5-fold molar excess over proteins. Labeling reactions were carried out for 3.5 h in the case of both proteins. Reactions were stopped by the addition of 50 mM Tris-HCl final concentration. The labeled proteins were purified with a PD10 (GE Healthcare) gel filtration column pre-equilibrated with Storage buffers (hSSB1: 25 mM Tris-HCl pH 8.0, 50 mM NaCl, 100 mM MgCl_2_, 10% glycerol, 1 mM DTT; INTS3: 25 mM Tris-HCl pH 8.0, 50 mM NaCl, 10% glycerol, 1 mM DTT). Samples were dialyzed against Storage buffers and in the case of INTS3 an Amicon Ultra 100K spin column (Sigma-Aldrich) was applied to remove residual free dye.

Protein concentrations were determined by Bradford method. Labeling efficiency was determined by visible light spectrometry using *ε*494 = 80000 for 5-IAF, ε488 = 73000 M^--1^ cm^-1^ for AF488, and ε647 = 270000 M^--1^ cm^-1^ for AF647. Labeling ratios were 60% for hSSB2, 17% for INIP, 10% for hSSB1, 51% for hOGG1, 55% for G3BP1, and 9.6% for INTS3.

Protein purity was checked for all constructs using SDS-PAGE. All constructs were frozen in liquid N2 in small aliquots and stored at -80°C. Labeling methods for BLM and EcSSB are described in (*23*).

### Fluorescence anisotropy titrations

DNA and RNA binding was measured in FP buffer (25 mM Tris-HCl pH 7.4, 50 mM KCl, 1 mM DTT) with 10 nM of 3’-fluorescein-labeled 36-mer ssDNA or 36-mer ssRNA oligonucleotide (ssDNA: ATTTTTGCGGATGGCTTAGAGCTTAATTGCGCAACG- fluorescein, ssRNA: AUUUUUGCGGAUGGCUUAGAGCUUAAUUGCGCAACG -fluorescein). Fluorescence anisotropy of 12-μl samples was measured in 384-well low-volume nontransparent microplates (Greiner Bio-one, PN:784900) at 25°C in a Synergy H4 Hybrid Multi-Mode Microplate Reader (BioTek) and converted to anisotropy values. Fits were performed using the Hill equation (*n* = 3 independent measurements).

### Turbidity measurements

Turbidity (light depletion at 600-nm wavelength) titrations were performed at indicated hSSB1 concentrations in the presence of indicated ssDNA (dT_18-96_ homopolymer, single-stranded deoxythymidine oligonucleotides) or ssRNA (U_32_, 32mer single-stranded uridine oligonucleotide) concentrations and were measured in a Tecan Infinite Nano+ plate reader instrument at 25°C. For measurements at low micromolar H_2_O_2_ concentrations (**Fig. 2E**), DTT was removed from hSSB1 storage buffer using a 10K spin column (Sigma-Aldrich) and DTT concentration was remeasured with DTNB (Ellman’s Reagent) to a final concentration of 0.4 µM.

### Epifluorescence microscopy for *in vitro* LLPS

A Nikon Eclipse Ti-E TIRF microscope was used in epifluorescence mode with apo TIRF 100x oil immersion objective (numerical aperture (NA) = 1.49). A Cyan 488-nm laser (Coherent), a 543-nm laser (25-LGP-193–230, Melles Griot) and a 642-nm laser (56RCS/S2799, Melles Griot) were used for excitation. Fluorescence was deflected to a ZT405/488/561/640rpc BS dichroic mirror and recorded by a Zyla sCMOS (ANDOR) camera. Images were captured using the imaging software NIS-Elements AR (Advanced Research) 4.50.00. Experiments were recorded with 2x2 binning and 200-ms laser exposure optical setup. 20-μl volumes of samples were introduced into μ-Slide Angiogenesis (Ibidi) microscope slides at 25°C. Sample components were mixed freshly and incubated for 1 h before imaging, unless indicated otherwise.

In the absence of nucleic acids (**Figs. 2A, S4E, S8B**) and for measurements shown in **Fig. 3C**, 5 µM hSSB1 and 0.1 µM Alexa Fluor 647-labeled hSSB1 (hSSB1^AF647^) were mixed. All nucleic acid-containing experiments were carried out using 5 µM hSSB1 (wild-type or variants) in the presence of 2 µM ssDNA (dT_79_, containing 100 nM Cy3-labeled dT_79_ or dT_45_, containing 100 nM Cy3-labeled dT_45_) or 2 µM ssRNA (U41; 41-mer uridine homopolymer, containing 100 nM Cy3-labeled nonhomopolymeric 41-mer ssRNA) unless indicated otherwise. **Fig. S2** shows that labeled ssDNA can readily enter hSSB1 droplets, thus it can be used for LLPS visualization without the need for protein labeling. For droplet fusion experiment, chambers were treated with 1.5 mg/ml Blocking Reagent (Roche) for 30 min before introducing hSSB1. Fusions were monitored at 4-5 sec intervals.

For multiprotein co-condensation experiments (**Fig. 5**), 180 nM labeled protein interaction partners were used either alone or together with 5 µM hSSB1 containing 0.1 µM AlexaFluor647-hSSB1. Each experiment contained also 2 µM dT_79_ (or 2 µM U_41_ in **Fig. S11B**) and 980 µM H_2_O_2_. In the case of labeled hSSB2, BLM, hOGG1, G3BP1, EGFP, and fluorescein dye molecules, hSSB1 was incubated for 1 hour before introducing fluorescent partners, then co-incubated with partners for additional 30 min before imaging. Since INTS3 and hSSB1 were both labeled with AlexaFluor647, in experiments containing labeled INTS3, labeled ssDNA (2 µM dT_79_ containing 0.1 µM Cy3-dT_79_) was used to visualize hSSB1 droplets (5 µM unlabeled protein). Furthermore, in experiments containing INTS3 and/or INIP partners, hSSB1 was co-incubated with partners for 2 hours before imaging. For the INIP + INTS3 complex-containing sample, 180 nM labeled INIP and 1 µM unlabeled INTS3 were used.

### Image processing for epifluorescence microscopy

ImageJ software was used to analyze unprocessed images. Image stack was generated from raw images of a given experiment to set brightness and contrast equally, using the automatic detection algorithm. Images were background corrected with the built-in rolling ball background correction, unless otherwise indicated. Montage was generated from stack of a given experiment to visually represent the changing conditions of the experimental set.

Since hSSB1 condensates spread out on the surface of microscope slide over time, instead of droplet size analysis (*23*) we analyzed the mean grey values and total areas of droplets. Stack of 3 images was generated for each condition. Middle ROI (Region of Interest) (area: 600 x 600 pixels; X,Y coordinates from left, uppermost pixel position: 300 pixels) of every stack was selected for technical reasons. Mean grey values (sum of intensity values from all pixels divided by the number of pixels) were measured for each middle ROI using the Stack Fitter plugin, then averaged for each condition separately. For determination of the total droplet area of middle ROIs, background correction thresholds were set using the built-in image thresholder of ImageJ. The particle analyzer algorithm (smallest detected size was set to 0.2 μm^2^, circularity 0.1–1) was applied to outline the distinct fluorescent spot areas of each middle ROI of a stack, which were added together separately and the distinct ROI values of one condition were averaged to get the total droplet area in μm^2^.

### *In vitro* oxidation experiments

20 μM of protein was incubated in LLPS buffer for 45 minutes at room temperature with indicated H_2_O_2_ concentrations. Covalent oligomers were separated via tris-glycine based SDS-PAGE on a 10% polyacrylamide gel. Electrophoresis was performed at 200 V constant voltage for 90 min. Gels were rinsed with ddH_2_O and stained overnight with PageBlue Protein Staining Solution (Thermo Fisher) and de-stained in ddH_2_O. Fraction of monomers and dimers were analyzed by pixel densitometry using the GelQuant Pro v12 software (DNR Bio Imaging Ltd.).

### Cell treatments, transfection, hSSB1 silencing

HeLa, HEK293T, and HFF-1 cells were maintained at 37°C under 5% CO_2_ atmosphere in RPMI-1640 medium (ThermoFisher, for HeLa) and DMEM (ThermoFisher, for HEK293T and HFF-1) supplemented with fetal bovine serum, gentamicin and amphotericin B.

For plasmid transfection, 25,000 cells were seeded onto plasma-sterilized, polylysine-treated coverslips in wells of a 24-well plate. Cells were transfected on the next day with pEGFP-C1 or pEGFP-C1-hSSB1 plasmids, encoding EGFP or EGFP-fused hSSB1 under the regulation of a CMV promoter. 1 µg of plasmid was transfected via lipofection using Lipofectamine 2000 reagent (ThermoFisher), in medium free of antibiotics and antifungals. Normal medium was replaced after 4 h. After 24 h of expression, cells were treated in serum free media as indicated.

For non-transfected cells, 25,000 cells were seeded into wells of a confocal chamber slide and treated the next day in serum free media as indicated. For hSSB1 silencing, 25,000 cells were seeded onto plasma-sterilized, polylysine-treated coverslips in wells of a 24-well plate. Cells were transfected on the next day with 25 nM ON-TARGETplus human NABP2 siRNA SMARTpool (L-014288-01-0005, Horizon) using DharmaFect transfection reagent, in a medium free of antibiotics and antifungals. For non-targeting RNA control, the ON-TARGETplus Non-targeting Control Pool (D-001810-10-05) was used. Normal medium was replaced after 6 h. 48 h after transfection, cells were lyzed for Western blot analysis or treated in serum free media as indicated.

### Cell survival assay

10,000 cells were seeded in wells of a 96-well plate in RPMI medium supplemented with 10% FBS. After two days cells were treated with H_2_O_2_ for 1 h. PrestoBlue reagent (Invitrogen) was added to a final concentration of 10% and was incubated for additional 1 h. PrestoBlue incubation time was taken into account for total treatment time. Fluorescence was measured at 590 nm with 560 nm excitation wavelength.

### Immunocytochemistry

Cells were washed with DPBS (Dulbecco’s Phosphate Buffered Saline) and fixed with 4 % PFA for 20 minutes. Membrane permeabilization was achieved with 0.5% Triton-X for 5 min. Samples were blocked with 2% BSA for 1 h at room temperature, and primary antibodies were applied overnight in blocking reagent at 4 °C. Anti-hSSB1 (HPA044615), anti-G3BP1 (ab56574), and anti-SK1-Hedls (sc-8418) were purchased from Sigma-Aldrich, Abcam, and Santa Cruz Biotechnology, respectively. Secondary antibodies (anti-mouse IgG AF647, anti-rabbit AF488, Thermo), conjugated with fluorophore were applied (1 h, room temperature) for fluorescence labeling. For experiments using coverslips coated with immunostained cells, immobilization was carried out using Mowiol 4.88 (Polysciences) (supplemented with DAPI) on microscope slides. Cells seeded onto confocal chamber slides were stained for chromatin using Hoechst 33342 or DAPI as indicated in the figures.

### Western blot

Cells were lyzed and suspended with 95°C 1x Laemmli-buffer (Bio-Rad) supplemented with 1 mM DTT. No DTT was added for non-reducing Western blot samples (**Fig. S13C-D**).

Samples were incubated at 95°C for 10 minutes and loaded onto SDS-polyacrylamide gels (Mini-Protean TGX gels 4-20%, Bio-Rad). Following gel electrophoresis, proteins were transferred onto nitrocellulose membranes. Blocking was achieved using TBST containing 5% BSA for 1 h at room temperature. Primary antibodies (anti-hSSB1 HPA044615 or anti-GAPDH G9545) were applied overnight in blocking reagent at 4°C. HRP-conjugated secondary antibodies (Peroxidase AffiniPure Goat Anti-Rabbit or Anti-Mouse IgG, Jackson ImmunoResearch) were applied for 1 h at room temperature. Immobilon Crescendo Western HRP substrate was used for chemiluminescence-based image development. Images were captured on a Bio-Rad ChemiDoc imaging system.

### Epifluorescence and confocal microscopy of immunostained cells

For epifluorescence microscopy, a Zeiss Cell Observer Z1 microscope was used with a plan-apochromat 63x oil immersion objective with a numerical aperture of 1.46. 12-bit images were captured at a pixel size of 1388 x 1040. LED module light sources were used for excitation at 385 nm, 475 nm, and 630 nm with excitation/emission filter setups of 335- 383/420-470, 450-490/500-550, 625-655/665-715 nm, respectively. An AxioCam MR R3 camera (Zeiss) was used for fluorescence detection.

For confocal microscopy, a Zeiss LSM 800 microscope was used with a plan-apochromat 63x oil immersion objective with a numerical aperture of 1.40. 16-bit images were obtained at 1437x1437 pixel size. For excitation, 405 nm, 488 nm, and 640 nm wavelength laser lights were used with pinhole sizes of 1 Airy unit in each case. GaAsP photomultiplier tubes were used for fluorescence emission detection. Imaging and 3D reconstruction were performed using ZENPro software.

### Colocalization analysis

Colocalization analysis was performed on confocal images using ImageJ. Mean intensity of G3BP1 inside SGs was measured and thresholds on the red channel were set as the minimum of the observed mean intensity of G3BP1 droplets. Thresholds on the green channel were set similarly, as the minimum of the observed mean intensity of hSSB1 inside SGs. Since hSSB1 has a high nuclear signal, nuclei were deleted from green channel images before thresholding. The thresholded images were used for calculating Manders split and object-based colocalization coefficients. Manders coefficient is proportional to the amount of fluorescence of the colocalizing pixels in each color channel. Values range from 0 to 1, expressing the fraction of intensity in a channel that is located in pixels where there is above-threshold intensity in the other color channel. Object-based colocalization analysis shows the ratios of centers of mass coincidence for particles detected in both channels using the thresholds described above. The analysis described above was performed using the JaCoP plugin in ImageJ (*61*).

### Determination of nuclear hSSB1 intensities and the number of SGs per cell

Using the microscopy setup described above, we obtained tile images at each different condition in order to achieve a sufficient sample size of around 60 cells per image. The tiles were stitched together using ZENPro. Nuclei were detected as ROIs based on DAPI/Hoechst signal. Integrated pixel density of hSSB1 signal was measured on nuclear ROIs and normalized to indicated controls.

SGs were counted on individual images according to G3BP1 signal, and SG counts were divided by the number of nuclei detected as ROIs in the given images based on DAPI/Hoechst staining. Thus, an average SG number per cell was determined from each individual image.

### Cell typization

To determine whether hSSB1 overexpression or silencing influences the fraction of cells forming stress granules (SG+), we devised a method to quantify the number of SG+ cells upon H_2_O_2_ treatment when either EGFP or hSSB1-GFP is overexpressed, or endogenous hSSB1 is silenced. Using the epifluorescence microscopy setup described above, we obtained tile images at each condition in order to achieve a sufficient sample size. Tiles were stitched together using ZENPro, and the full image was equally divided into 4 subimages, which were analyzed separately. Outlines of cells were detected based on G3BP1 signal using Cellpose deep learning-based algorithm (*62*). The detected outlines were imported to ImageJ as ROIs. Thus, each ROI represented a different cell in the image. Mean intensity, Skewness and Kurtosis (third and fourth order moment about the mean) were measured on ROIs. A cell was considered as EGFP+ if its mean fluorescence intensity in the green channel was higher than the mean intensity of the predetermined green autofluorescence of a HeLa cell in the applied imaging setup. Skewness and Kurtosis was indicative of the heterogenous distribution of the G3BP1 fluorescence signal in the cytoplasm. The more SGs were seen, the higher Skewness and Kurtosis values were obtained. Thus, a cell was considered SG+ if both Skewness and Kurtosis were higher than their predetermined thresholds of 1 and 5, respectively. Cell typization was done on all 4 subimages (*n* = 4) at the given condition. Means ± SEM (standard error of mean) values are reported.

### Quantification and statistical analysis

Data analysis and visualization was performed using OriginLab 8.0 (Microcal corp.). Pixel densitometry in electrophoretograms and immunoblots was performed using GelQuant Pro software v12 (DNR Bio Imaging Ltd.). Statistical analysis was performed in OriginLab 8.0. Significance levels are indicated in the figure legends.

## ACKNOWLEDGMENTS

### Funding

This work was supported by the “Momentum” Program of the Hungarian Academy of Sciences (LP2011-006/2011), ELTE KMOP-4.2.1/B-10-2011-0002, NKFIH K-123989, and NKFIH K-134595 grants to M.K. The project was supported by the NRDIO (VEKOP- 2.3.3-15-2016-00007 to ELTE) grant. G.M.H. was supported by the Premium Postdoctoral Program of the Hungarian Academy of Sciences (PREMIUM-2017-17 to G.M.H.). Z.J.K. and J.P. were supported by the New National Excellence Program of the Ministry for Innovation and Technology (Grants ÚNKP-21-3 to Z.J.K. and ÚNKP-19-2 to J.P.), and the Co-operative Doctoral Program of the Ministry of Innovation and Technology financed from the National Research, Development and Innovation Fund. This work was completed in the ELTE “SzintPlusz” Thematic Excellence Programme supported by the Hungarian Ministry for Innovation and Technology, and also in the framework of Project no. 2018-1.2.1-NKP-2018- 00005 implemented with the support provided from the National Research, Development and Innovation Fund of Hungary, financed under the 2018-1.2.1-NKP funding scheme.

### Author contributions

GMH, JP, ZJK, BJ, GS, and MK conceptualized the project and designed experiments. JP, ZJK, BJ, GMH, KT, HHP, JH, LM, NK, and ST performed experiments and analyzed the data. JP, GMH, ZJK, and KM wrote the paper. All authors participated in revising the manuscript.

### Data and materials availability

HFF-1 cell line was a gift by Ágota Apáti (HUN-REN Hungarian Research Network, Institute of Molecular Life Science). PDB files 4MZ9, 5D8E, 5D8F and 4OWW were obtained from the Protein Data Bank. hSSB1 expression levels were obtained from Expression Atlas (https://www.ebi.ac.uk/gxa/home). All data needed to evaluate the conclusions in the paper are present in the paper and/or the Supplementary Materials.

