## Supplementary Material for "Redox-dependent condensation and cytoplasmic granulation by human ssDNA binding protein 1 delineate roles in oxidative stress response"

[1] ELTE-MTA "Momentum" Motor Enzymology Research Group, Department of Biochemistry, Eötvös Loránd University, Pázmány P. s. 1/c, H-1117 Budapest, Hungary
[2] Present address: Laboratory of Single Molecule Biophysics, National Heart, Lung and Blood Institute, National Institutes of Health, Bethesda, Maryland 20892, USA
[3] HUN-REN-ELTE Motor Pharmacology Research Group, Department of Biochemistry, Eötvös Loránd University, Pázmány P. s. 1/c, H-1117 Budapest, Hungary
[4] Department of Physiology and Neurobiology, Eötvös Loránd University, Pázmány P. s. 1/c, H-1117 Budapest, Hungary
[5] HUN-REN Institute of Molecular Life Sciences, Research Centre for Natural Sciences, Magyar Tudósok körútja 2, H-1117 Budapest, Hungary
[6] Center for Cancer Research, Medical University of Vienna, Borschkegasse 8a, 1090 Wien, Austria
† Authors contributed equally.

**Supplementary Figures**

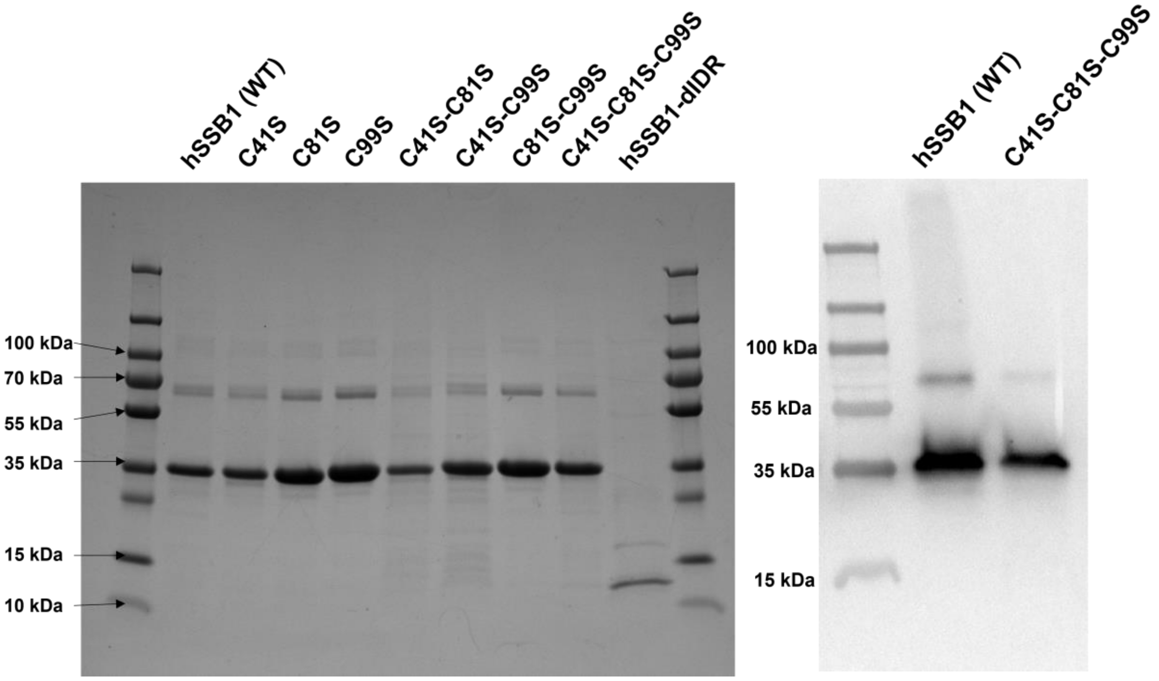


**Fig. S1. SDS-PAGE Coomassie-stained electrophoretogram (left) and immunoblot (right) of hSSB1 constructs in reducing conditions.** Left: Coomassie-stained electrophoretogram of purified hSSB1 (WT, wild type), its Cys-to-Ser substituted variants, and hSSB1-dIDR on a 12 % polyacrylamide gel in reducing conditions. We note that the mobility of hSSB1 constructs (except for hSSB1-dIDR) is lower than expected based on their molecular weight of 22 kDa, which is a frequently observed feature for proteins containing long ID regions. The mobility of hSSB1-dIDR, due to the lack of an ID region, corresponds to its actual molecular weight of 12 kDa. 1 μg protein was loaded, except for hSSB1-dIDR (0.5 μg). Right: Immunoblot of reduced hSSB1 WT and the C41S-C81S-C99S variant from a 4-20 % polyacrylamide gel. The protein band around 70 kDa, visible on both the Coomassie-stained electrophoretogram and the immunoblot, appears to be an SDS and DTT resistant non-covalent dimeric species. The presence of this species, however, did not affect the determination of the extent of oxidation-induced covalent dimerization of hSSB1 (**Fig. 4, S8**).

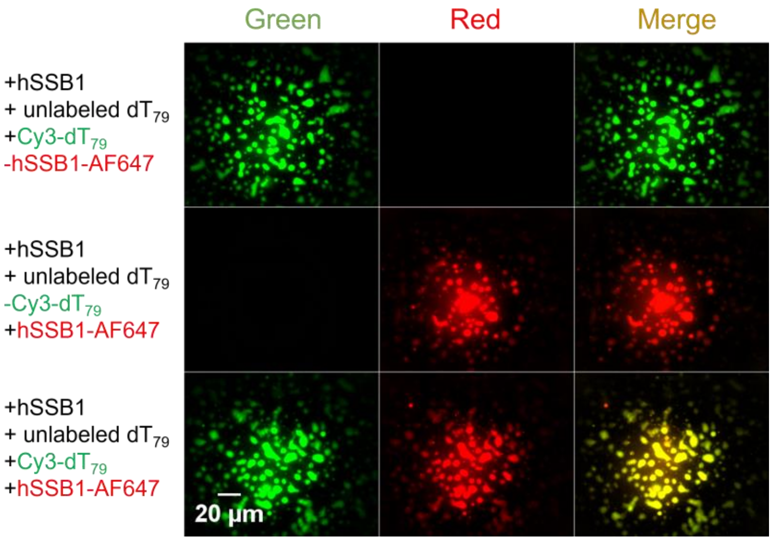


**Fig. S2. ssDNA is readily enriched inside hSSB1 droplets and can thus be used for hSSB1 LLPS visualization.** Top row shows hSSB1 (5 µM) droplets in the presence of labeled ssDNA (2 µM $dT_{79}$ containing 0.1 µM Cy3-$dT_{79}$). Middle row shows labeled hSSB1 (5 µM hSSB1 containing 0.1 µM hSSB1-AF647 (AlexaFluor647-labeled hSSB1)) droplets in the presence of unlabeled ssDNA (2 µM $dT_{79}$). Bottom row shows labeled hSSB1 (5 µM hSSB1 containing 0.1 µM hSSB1-AF647) droplets in the presence of labeled ssDNA (2 µM $dT_{79}$ containing 0.1 µM Cy3-$dT_{79}$). 980 µM $H_2O_2$ was used to initiate LLPS. Green channel shows Cy3-labeled $dT_{79}$, red channel shows hSSB1-AF647, yellow color indicates co-condensation.

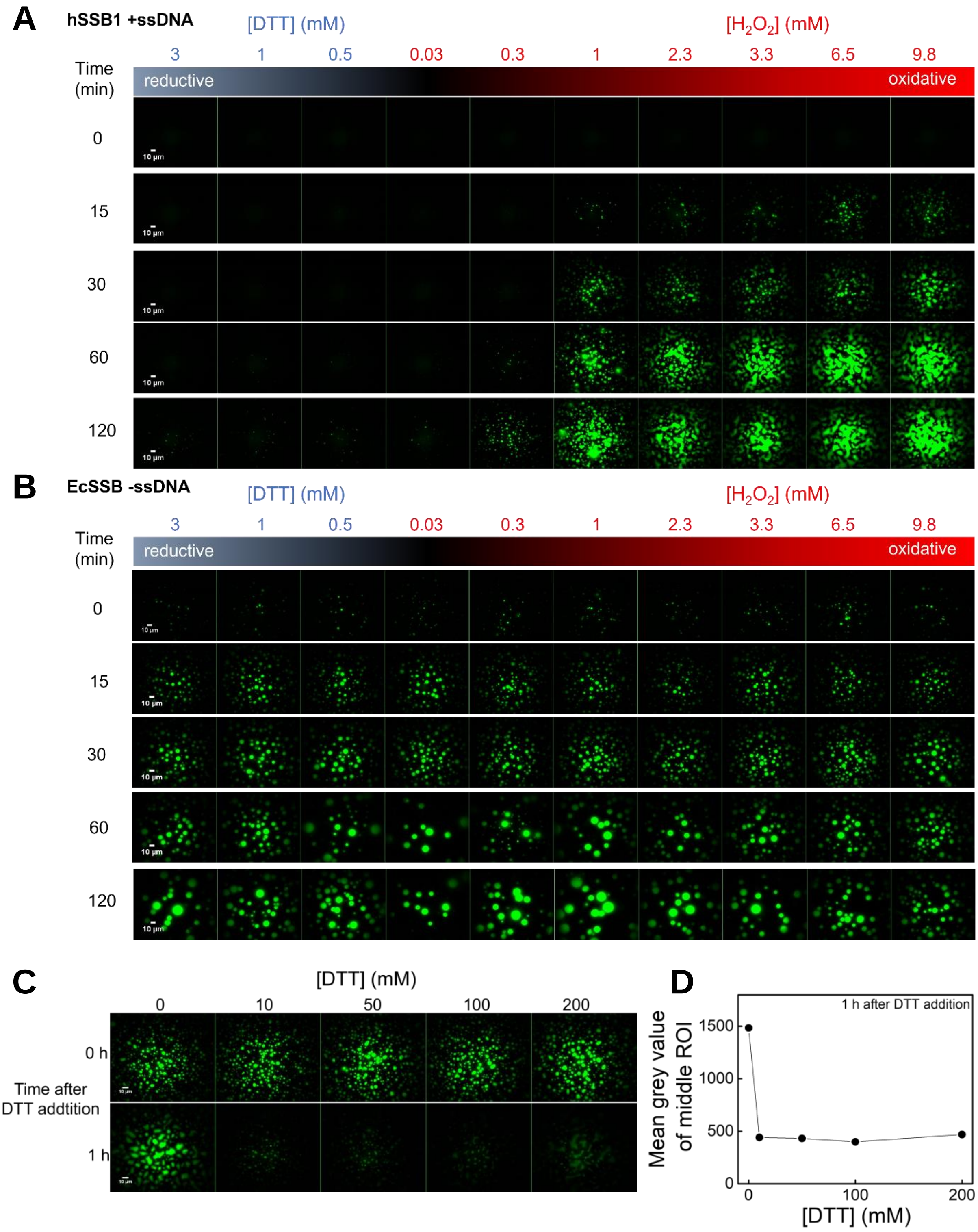


**Fig. S3. hSSB1 droplets form selectively and reversibly under oxidative conditions. (A)** Time-lapse epifluorescence microscopy images of the dependence of hSSB1 (5 μM) droplet formation on redox conditions, in the presence of labeled ssDNA (2 μM $dT_{79}$ containing 0.1 μM Cy3-$dT_{79}$) Droplets formed selectively in oxidizing conditions in less than 15 minutes. **(B)** Control experiments for panel **A** performed using EcSSB, showing that EcSSB droplet formation is unaffected by redox conditions. Condensates were visualized via labeled EcSSB

(9 µM EcSSB containing 0.15 µM AlexaFluor 555-labeled EcSSB). **(C)** Droplet formation by hSSB1 is reversible, as droplets dissolve upon addition of reducing agent. Top row shows epifluorescence microscopy images of samples containing hSSB1 (5 µM) in the presence of 2 µM $dT_{79}$ (containing 0.1 µM Cy3-$dT_{79}$) in LLPS buffer supplemented with 980 µM $H_2O_2$. Samples were incubated for 1 hour before the addition of indicated concentrations of DTT, and then incubated for additional 1 hour (bottom row). **(D)** Mean grey value of middle ROIs of each image in experiments shown in the bottom row of panel **C**.

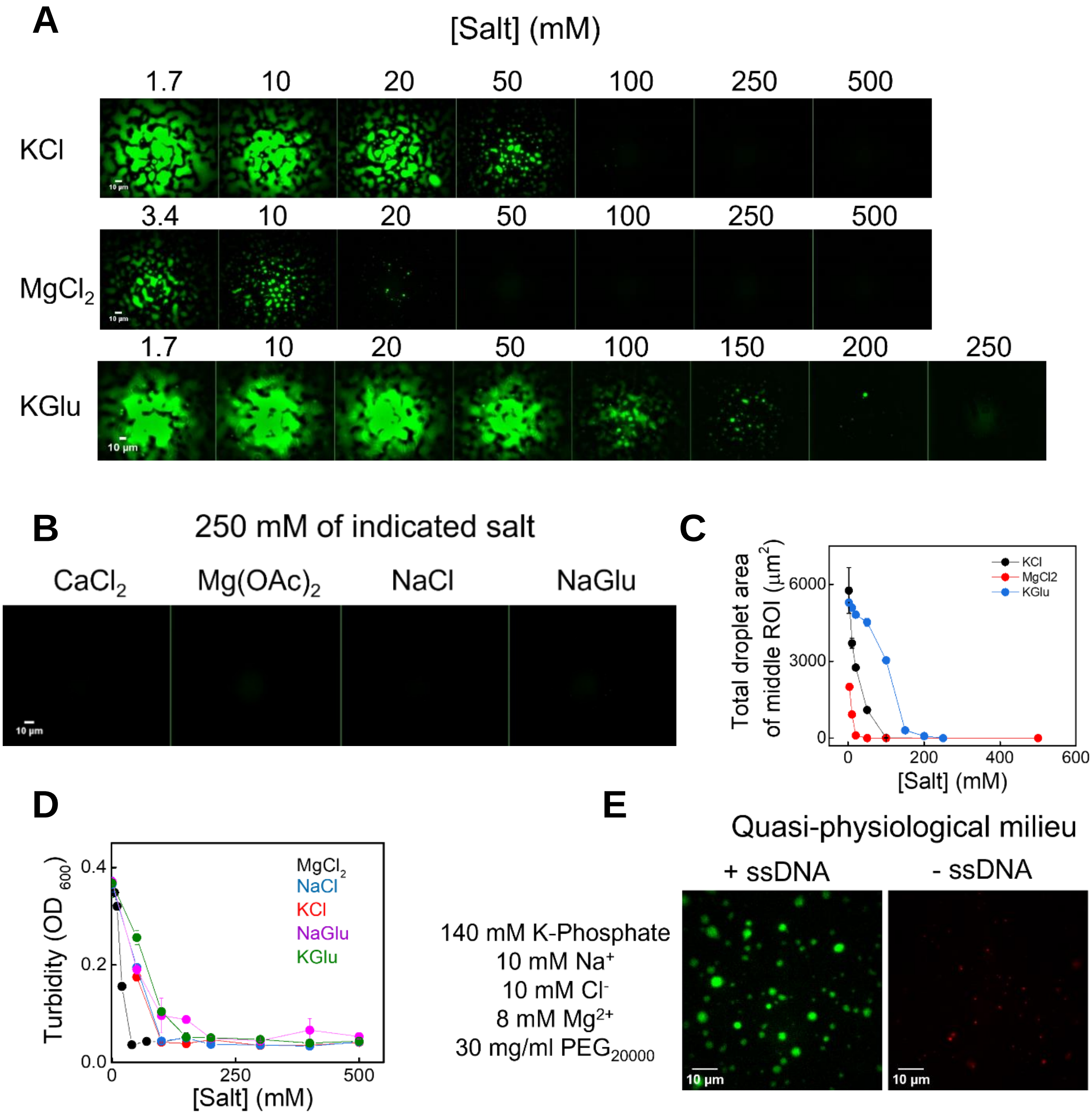


**Fig. S4. Ion concentration dependence of hSSB1 phase separation. (A)** Epifluorescence microscopy images of samples containing hSSB1 (5 µM) in the presence of labeled ssDNA (2 µM $dT_{79}$ containing 0.1 µM Cy3-$dT_{79}$) in no salt LLPS buffer (see Methods) supplemented with the indicated salt concentrations. Droplet formation was initiated with 980 µM $H_2O_2$. Experiments using KCl and KGlu (potassium glutamate) contained 10 mM $MgCl_2$, and magnesium titration experiments contained 50 mM KCl, due to technical reasons. **(B)** Effects of additional types of salts on hSSB1 LLPS. Fluorescence microscopy images are shown for hSSB1 (5 µM) in the presence of $dT_{79}$ (2 µM $dT_{79}$ containing 0.1 µM Cy3-$dT_{79}$) in no salt LLPS buffer containing 250 mM of the indicated salt. Droplet formation was initiated by adding 980 µM $H_2O_2$. **(C)** Total droplet area analysis of middle ROIs of 3 images from the experiment in panel **A**. Mean ± SEM values are shown. **(D)** Turbidity measurements (5 µM hSSB1, 2 µM $dT_{79}$, 980 µM $H_2O_2$ in no salt LLPS buffer) also confirmed the inhibitory effects of supraphysiological salt concentrations. Means ± SEM are shown for $n$ = 3 independent measurements. **(E)** In a phosphate buffered (pH 7.5), quasi-physiological milieu of 140 mM $K^+$, 10 mM NaCl, 8 mM $Mg(OAc)_2$, 30 mg/ml $PEG_{20000}$, hSSB1 was able to undergo LLPS in the presence of ssDNA (left: 2 µM $dT_{79}$ containing 0.1 µM Cy3-$dT_{79}$ together with 5 µM hSSB1, right: 5 µM hSSB1 containing 0.1 µM hSSB-AF647) in response to $H_2O_2$ (980 µM) treatment.

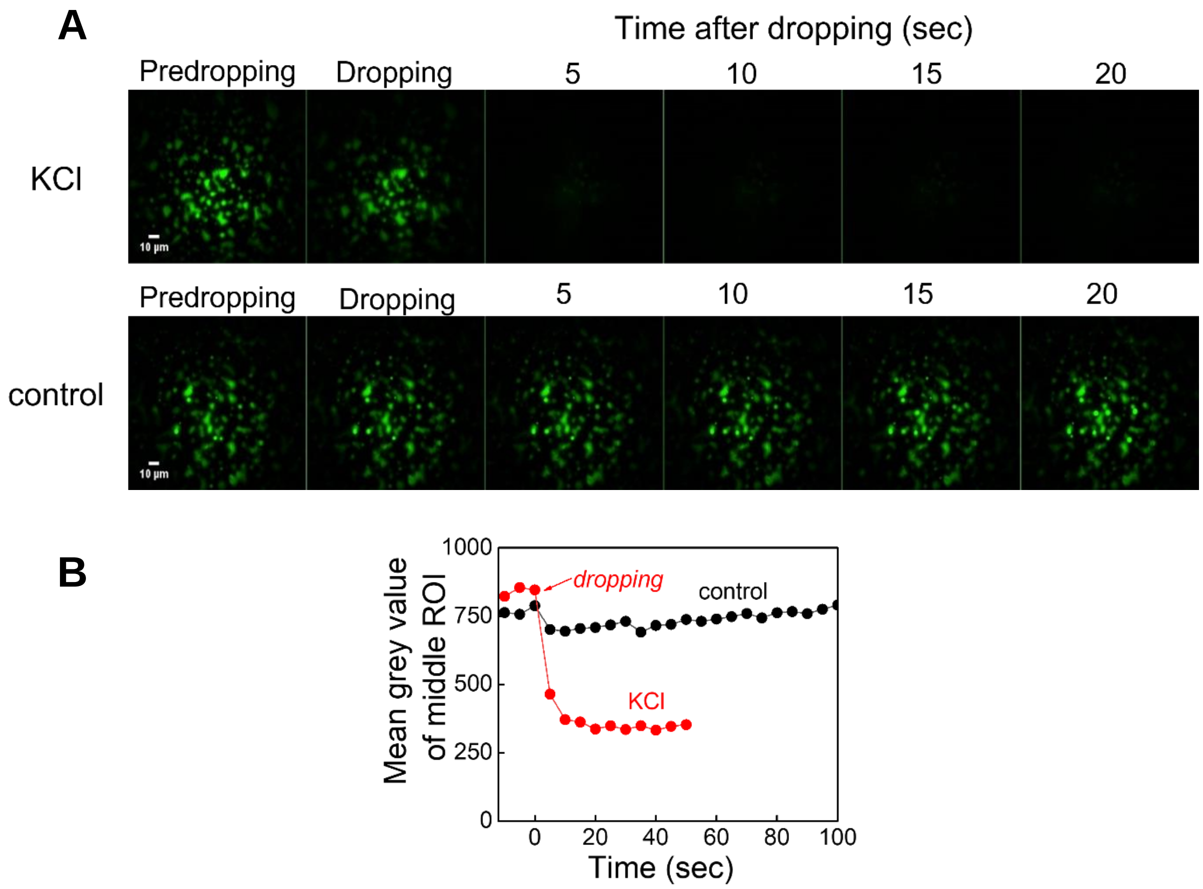


**Fig. S5. Rapid increase in ion concentration result in the dissolution of hSSB1 droplets, confirming reversibility of droplet formation. (A)** hSSB1 droplets dissolve upon addition of supraphysiological concentration of KCl (500 mM final). Epifluorescence microscopy images of samples containing hSSB1 (5 µM) in the presence of labeled ssDNA (2 µM $dT_{79}$ containing 0.1 µM Cy3-$dT_{79}$) in LLPS buffer supplemented with 980 µM $H_2O_2$. Samples were incubated for 1 hour before the addition of KCl (500 mM final concentration, top row) or Tris buffer (25 mM pH 7.4, bottom row) as control, and then visualized in 5-second intervals. **(B)** Mean grey value of middle ROIs of each image in experiments shown in panel **A.**

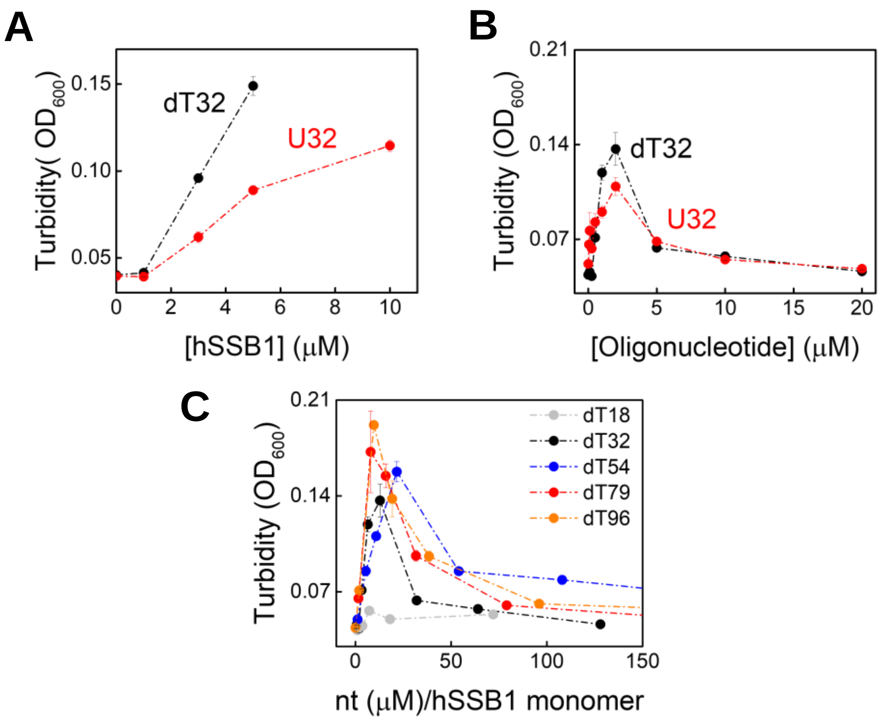


**Fig S6. ssDNA and ssRNA exert similar effects on hSSB1 condensation.**
(**A**) Turbidity ($OD_{600}$) of samples in which either ssDNA (2 μM dT32) or ssRNA (2 μM U32) were titrated with hSSB1 in the presence of $H_2O_2$ (980 μM). (**B**) Turbidity ($OD_{600}$) of samples in which hSSB1 (5 μM) was titrated with ssDNA (dT32) or ssRNA (U32) in the presence of $H_2O_2$ (980 μM). (**C**) Turbidity ($OD_{600}$) of samples in which hSSB1 (5 μM) was titrated with dT homopolymers of different length in the presence of $H_2O_2$ (980 μM). Maximal turbidity values correspond to an ssDNA interaction stoichiometry of around 10-15 nucleotides (nt) / hSSB1 monomer. In all panels, error bars represent SEM for $n$ = 3 independent measurements.

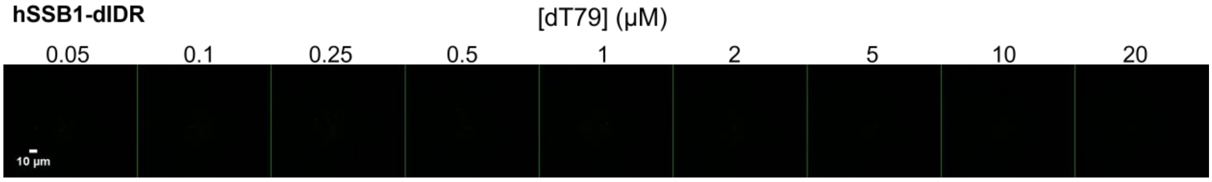


**Fig. S7. hSSB1-dIDR is unable to undergo LLPS, regardless of ssDNA interaction.** Epifluorescence microscopy images of samples containing hSSB1-dIDR (5 µM) in the presence of indicated ssDNA ($dT_{79}$) concentrations. 0.1 µM Cy3-$dT_{79}$ was present in each condition in LLPS buffer supplemented with 980 µM $H_2O_2$.

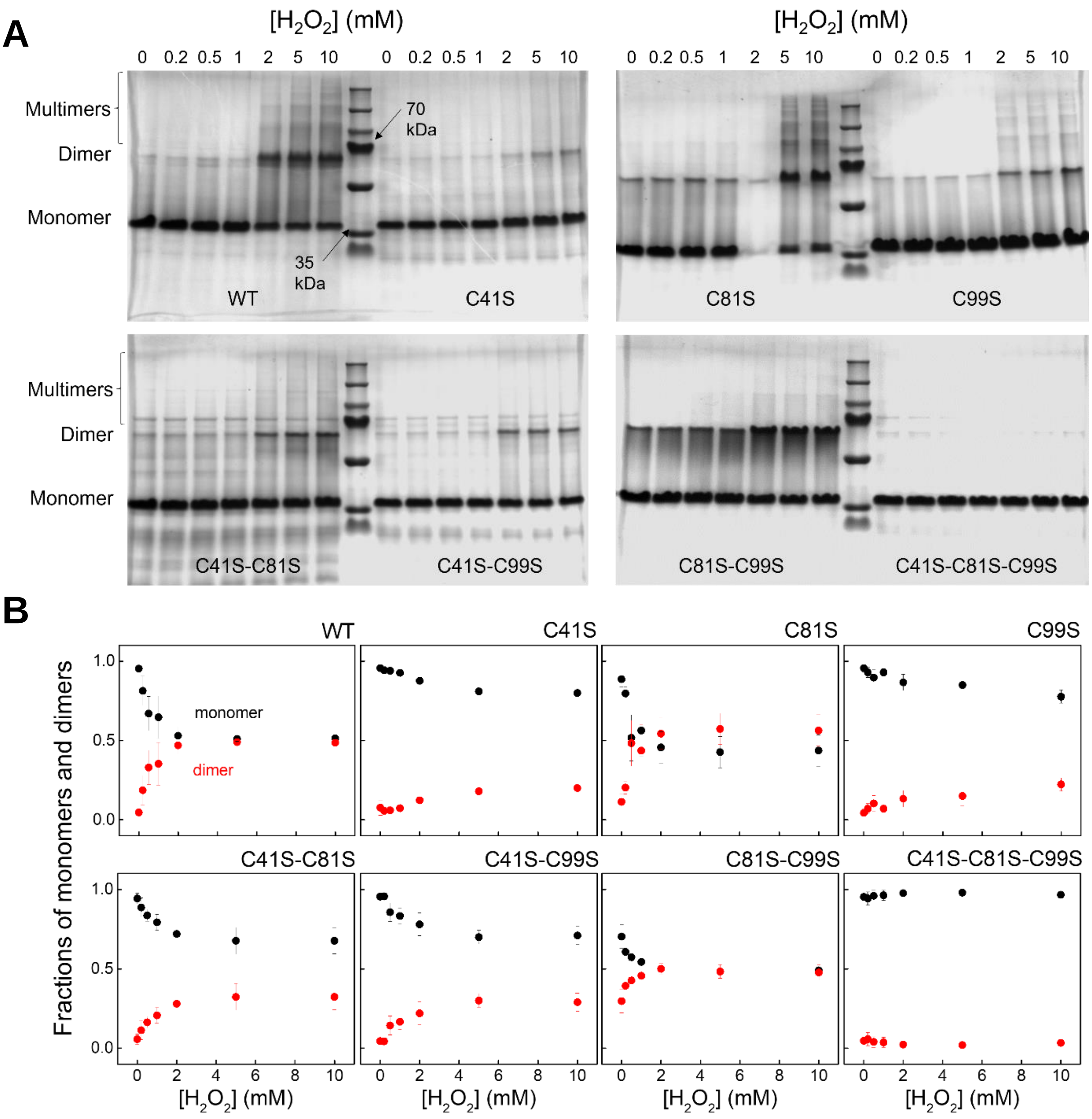


**Fig. S8. Effect of hSSB1 cysteine-to-serine substitutions on covalent oligomerization. (A)** Representative SDS-PAGE electrophoretograms (*n* = 3 independent measurements) of oxidation experiments for hSSB1 variants (cf. **Fig. S1**). Densitometric analysis was applied to determine changes in monomer : dimer ratios. Higher-order oligomers were omitted from analysis (see Materials and Methods). **(B)** Densitometric analysis of hSSB1 variants shows changes in monomer : dimer ratio in response to $H_2O_2$ treatment. Means ± SEM are shown for *n* = 3 independent measurements.

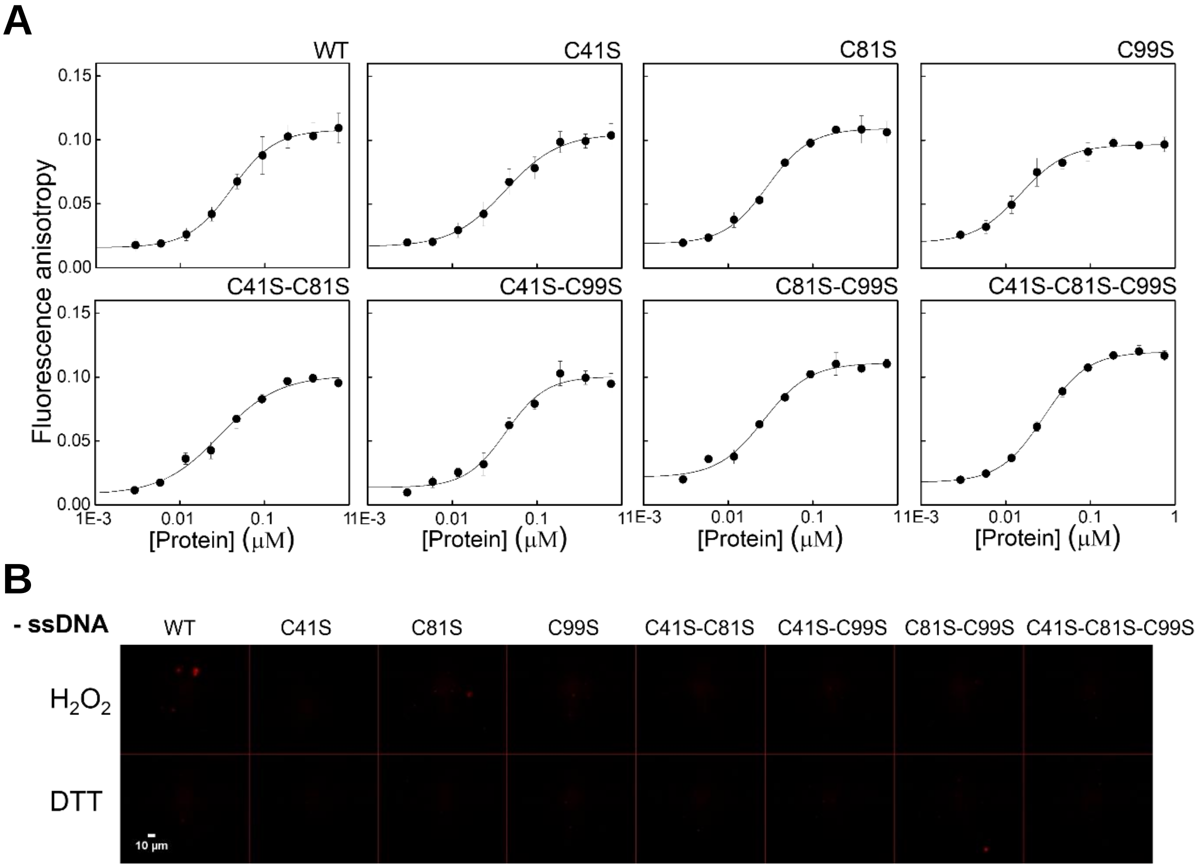


**Fig. S9. ssDNA binding by hSSB1 is largely unaffected by cysteine-to-serine substitutions. (A)** Results are shown for fluorescence anisotropy titrations using 10 nM ssDNA (3'-fluorescein-labeled 36-mer, see Materials and Methods). Solid lines show fits using the Hill-equation. Means ± SEM are shown for $n$ = 3 independent experiments in each case. Determined equilibrium dissociation constants ($K_d$) and Hill coefficients ($n$) are shown in **Table S1**. **(B)** hSSB1 variants are unable to undergo LLPS in the absence of ssDNA in either reducing or oxidizing conditions. Epifluorescence microscopy images are shown of indicated proteins (5 µM SSB variant together with 0.1 µM AF647-labeled wild type hSSB1) in LLPS buffer containing 1 mM DTT/980 µM $H_2O_2$.

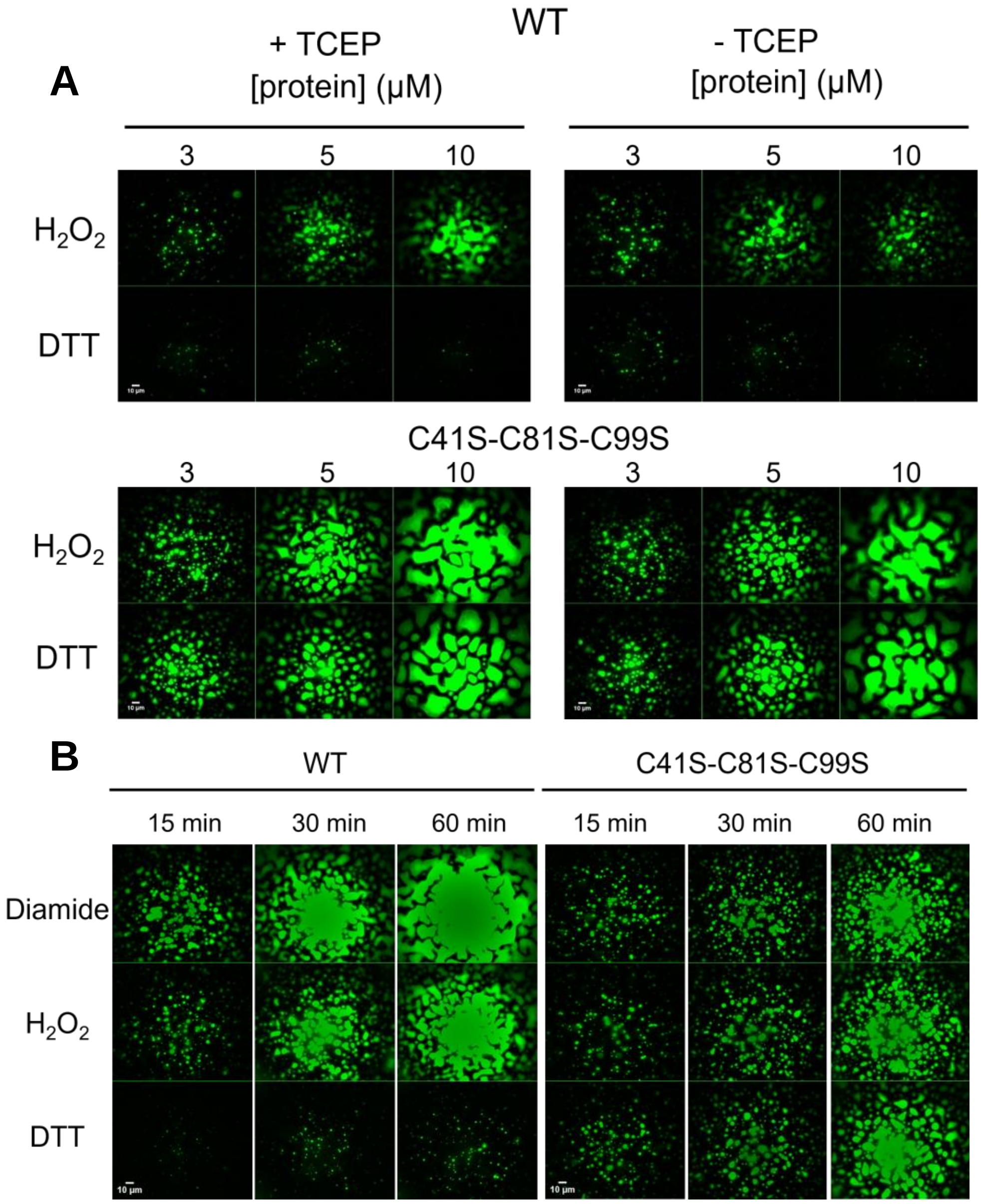


**Fig S10. Covalent oxidation of cysteines directly contributes to redox-sensitive LLPS of hSSB1.**

(**A**) Epifluorescence microscopic images of LLPS condensates formed by 5 μM hSSB1 WT (top) and C41S-C81S-C99S (bottom) in the presence of 2 μM dT79 (containing 0.1 μM Cy3-dT79) in LLPS buffer. Left panel shows condensation after overnight reduction by TCEP followed by buffer exchange. Right panel shows condensation by protein samples not treated with TCEP. TCEP treatment did not affect LLPS propensity and redox sensitivity. This result implies that the loss of redox-sensitive LLPS by C41S-C81S-C99S (and other variants, see **Fig. 4E**) is not a result of protein oxidation during storage. (**B**) Epifluorescence time-lapse images of LLPS condensates formed by 5 μM hSSB1 WT (left) and C41S-C81S-C99S (right) in the presence of 2 μM dT79 (containing 0.1 μM Cy3-dT79) in response to DTT, $H_2O_2$, and diamide (1 mM in each case). Diamide induces similar droplet formation as does $H_2O_2$ for the WT protein, while C41S-C81S-C99S is unaffected by redox conditions and diamide, indicating the role of cysteine oxidation in redox dependent LLPS.

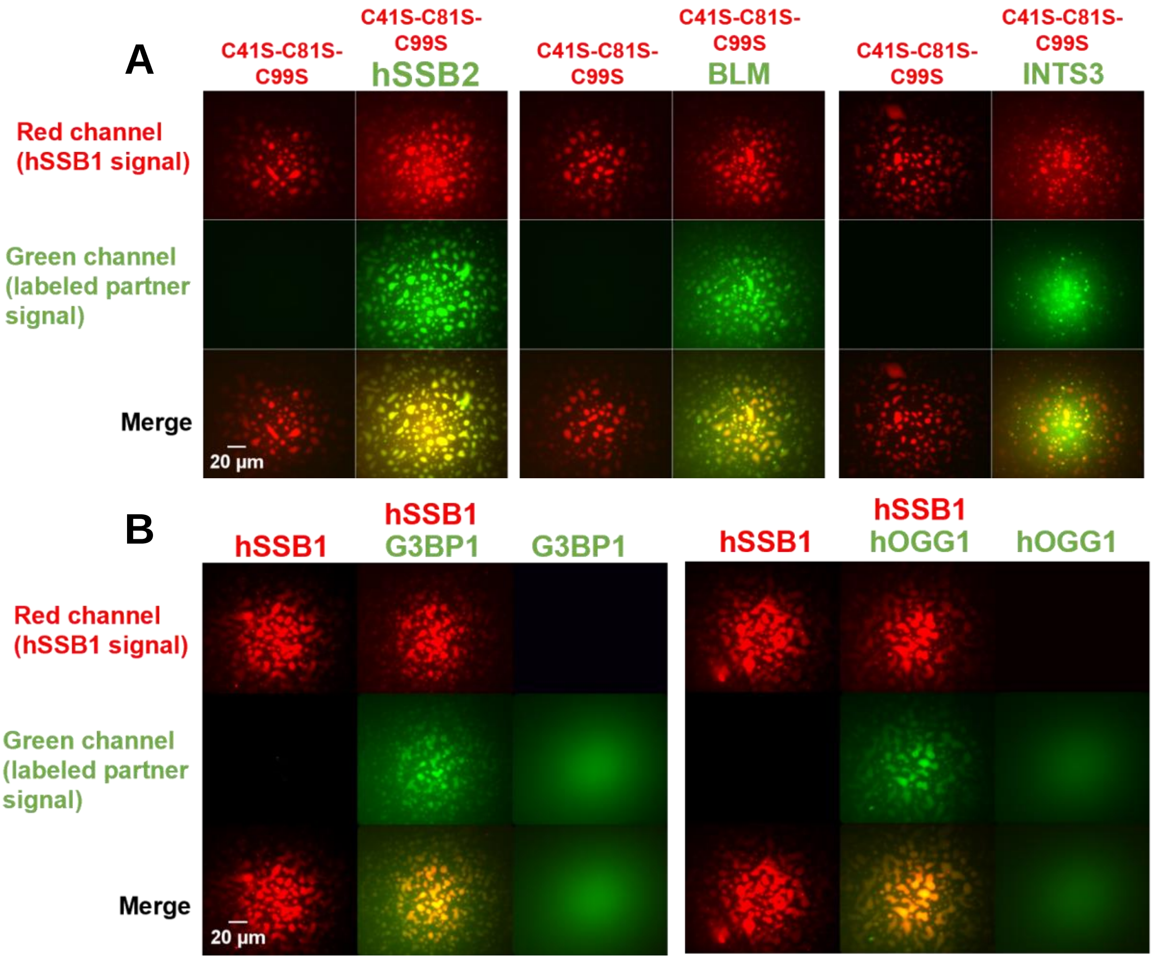


**Fig S11. Enrichment of genome maintenance proteins inside hSSB1 droplets is not a result of disulfide-crosslinking; G3BP1 and hOGG1 are enriched in hSSB1 droplets containing ssRNA**

(**A**) Two-channel fluorescence microscopic images showing enrichment of proteins inside hSSB1 C41S-C81S-C99S droplets. Columns represent two separate experiments (C41S-C81S-C99S, C41S-C81S-C99S + partner), while rows represent fluorescence channels. 5 µM C41S-C81S-C99S with 0.1 µM AlexaFluor647-labeled hSSB1 WT were present in the samples, together with ssDNA (2 µM dT79) and $H_2O_2$ (980 µM). 180 nM labeled interaction partner was used. In experiments containing labeled INTS3, labeled ssDNA (2 µM dT79 containing 0.1 µM Cy3-dT79) was used to visualize hSSB1 droplets (5 µM unlabeled protein). See Materials and Methods for further details. Red channel shows C41S-C81S-C99S droplets, green channel shows labeled interaction partners. Yellow color indicates co-condensation. Images were not background corrected. (**B**) Two-channel fluorescence microscopic images showing enrichment of G3BP1 and hOGG1 inside hSSB1 droplets containing ssRNA instead of ssDNA. Columns represent three separate experiments (hSSB1, hSSB1 + partner, partner alone), while rows represent fluorescence channels. 5 µM hSSB1 with 0.1 µM AlexaFluor647-labeled hSSB1 were present in the samples, together with ssRNA (2 µM $U_{41}$) and $H_2O_2$ (980 µM). 180 nM labeled interaction partner was used. Red channel shows hSSB1 droplets, green channel shows labeled interaction partners. Yellow color indicates co-condensation. Images were not background corrected.

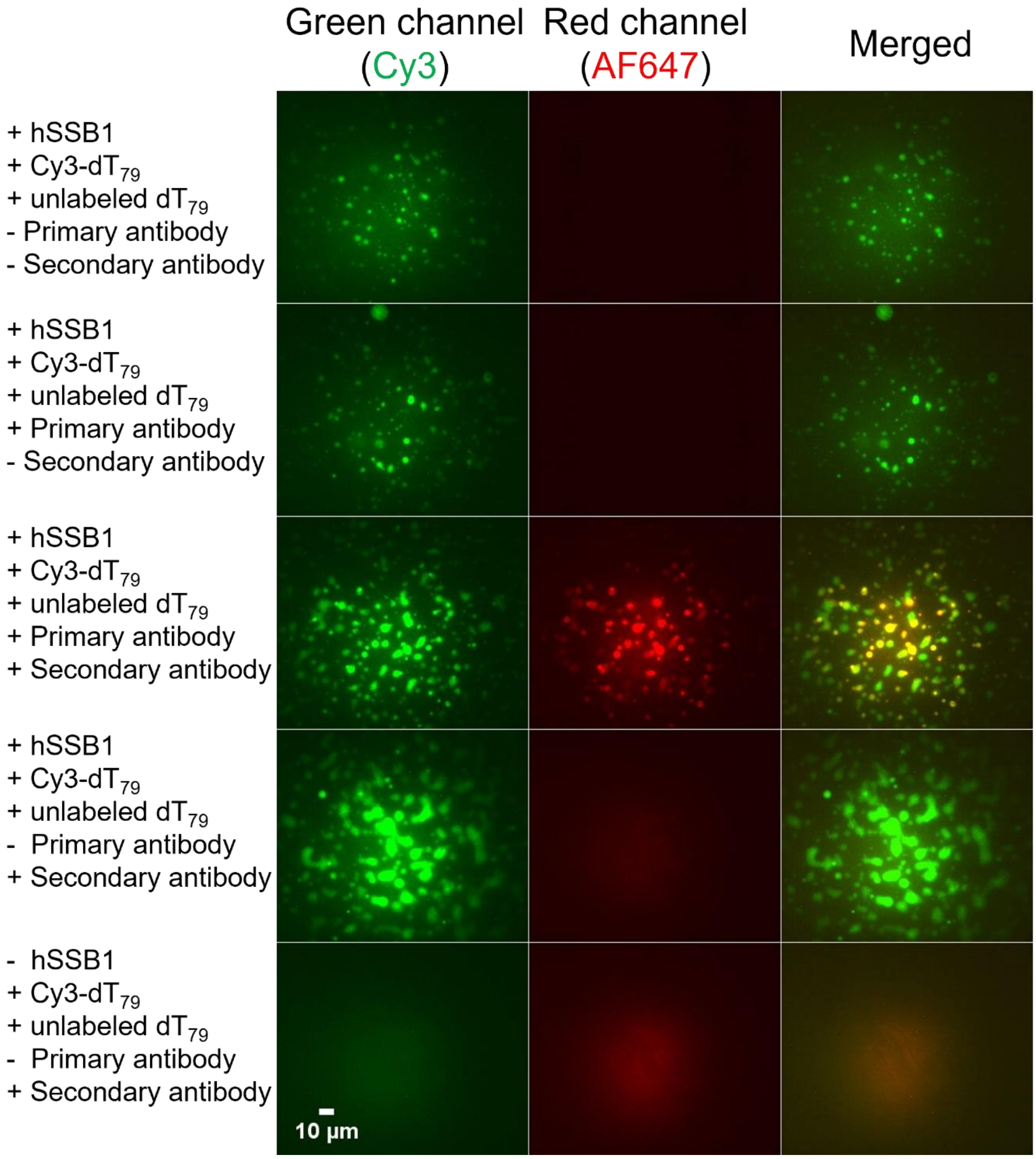


**Fig. S12. Anti-hSSB1 antibody can enter and stain hSSB1 droplets, enabling immunostaining of hSSB1 under LLPS conditions.** Epifluorescence microscopy images of samples containing hSSB1 (5 μM) in the presence of labeled ssDNA (2 μM $dT_{79}$ containing 0.1 μM Cy3-$dT_{79}$) in LLPS buffer supplemented with 980 μM $H_2O_2$. Anti-hSSB1 primary antibody (see Materials and Methods) was visualized through interaction with AlexaFluor647-labeled secondary antibody. Green channel shows hSSB1 droplets (*via* Cy3-$dT_{79}$ signal), red channel shows immunostaining. Yellow color indicates co-condensation. Droplets are only visible in the red channel when both primary and secondary antibodies are present, confirming the suitability of immunostaining of hSSB1 droplets.

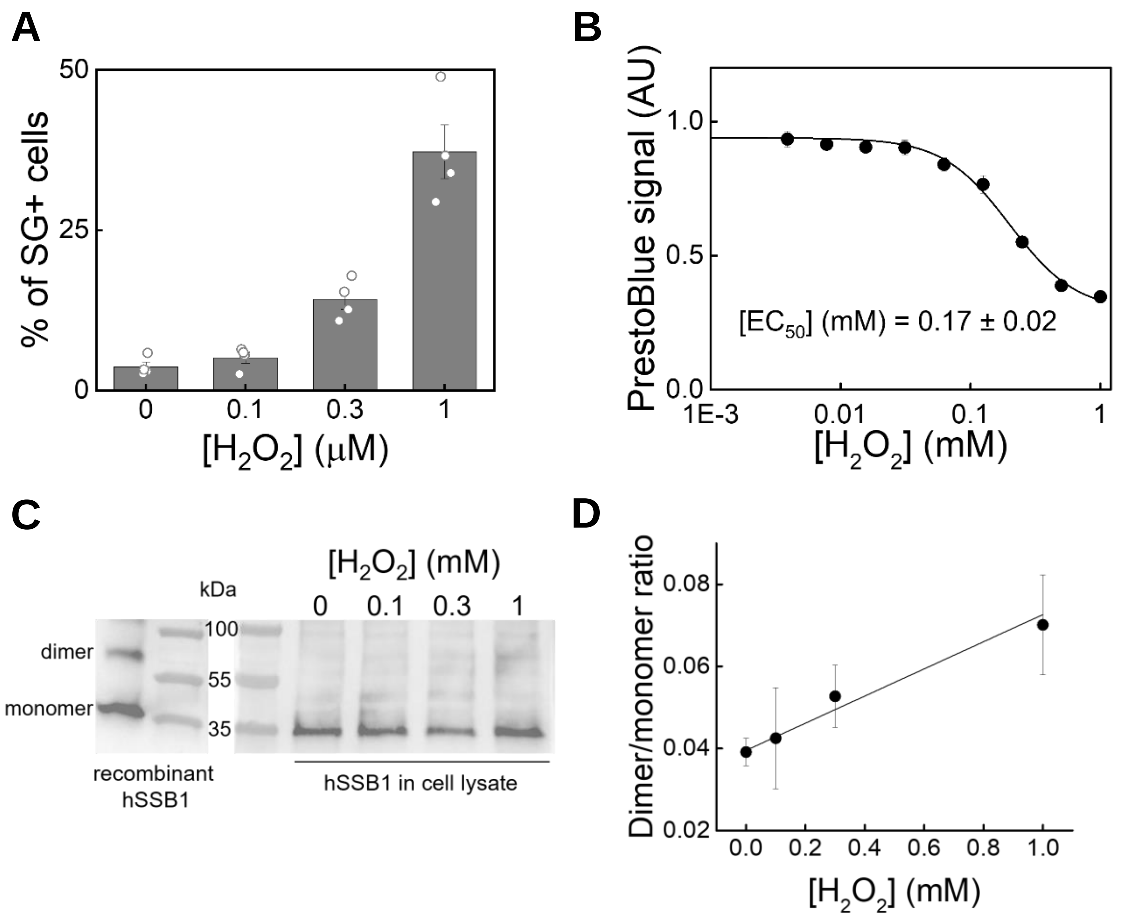


**Fig. S13. $H_2O_2$ treatment induces stress granule formation and decreases cell viability, while hSSB1 is partially oxidized into dimers in HeLa cells.** (**A**) Results of immunocytochemistry-based cell typization (see Materials and Methods) showing $H_2O_2$-induced SG formation. Means ± SEM are shown together with individual data points. (**B**) PrestoBlue-based cell survival curves of HeLa cells. Solid line shows fits based on the Hill equation. Means ± SEM are shown for $n = 3$ independent experiments. (**C**) Non-reducing immunoblots of recombinant hSSB1 (left) and of lysates from cells treated with indicated concentrations of $H_2O_2$ for 2 h (right). (**D**) An increase in dimer/monomer ratio of cellular hSSB1 is seen upon $H_2O_2$ treatment. Pixel densitometry data were obtained from immunoblots represented in panel **C**. Means ± SEM are shown for $n = 5$ independent measurements. Solid line shows linear fit (slope: 0.033 ± 0.004 $mM^{-1}$, intercept: 0.040 ± 0.001). Slope is significantly different from zero ($p < 0.05$).

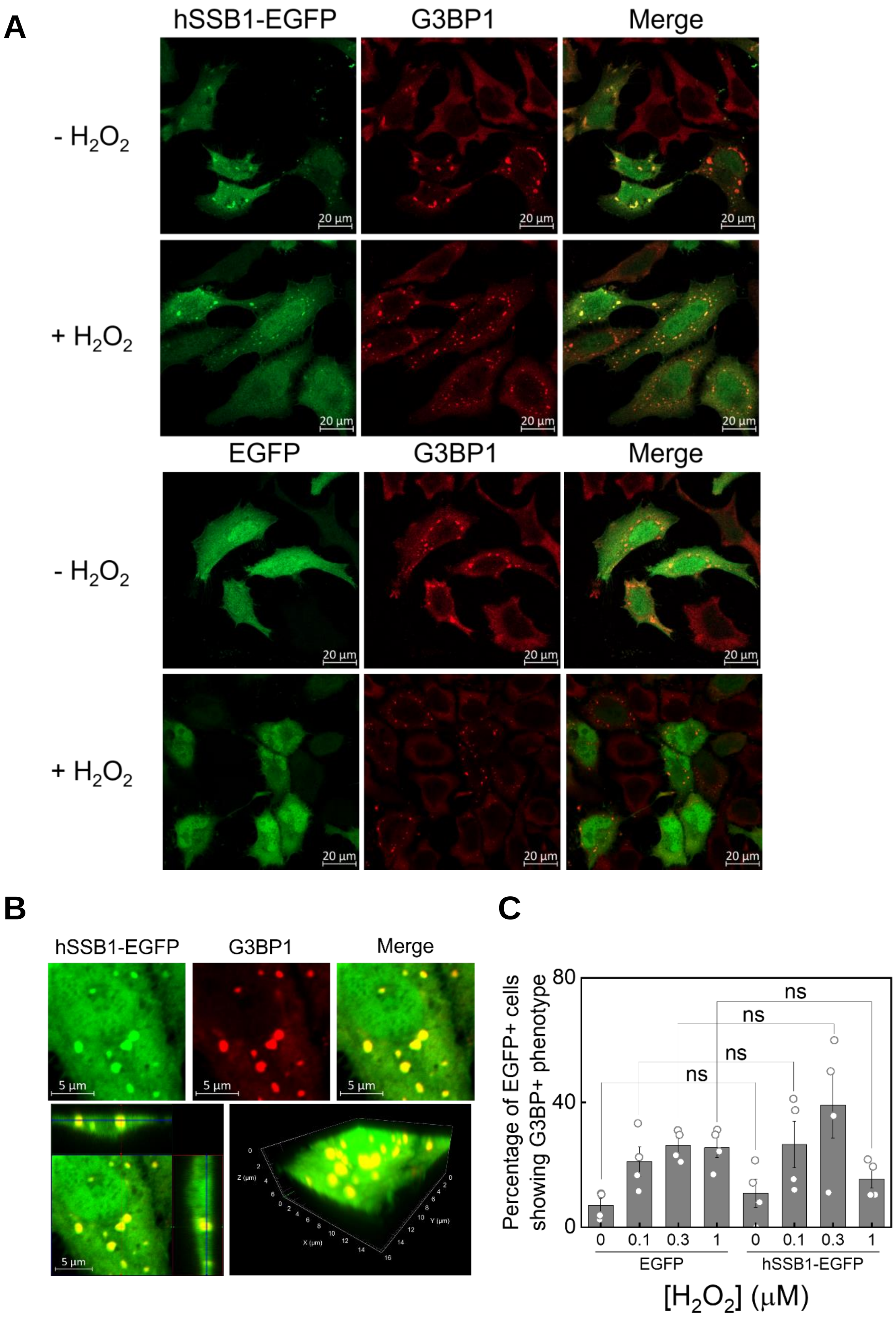


**Fig. S14. Transiently overexpressed hSSB1-EGFP colocalizes with stress granules. (A)** Representative confocal microscopy images ($n$ = 3 independent measurements) of HeLa cells transiently overexpressing EGFP-fused hSSB1 or EGFP alone. Top row shows untreated cells, bottom row shows cells treated with 0.3 mM $H_2O_2$ for 2 h. Green channel shows hSSB1-EGFP or EGFP, red channel shows immunofluorescence of G3BP1 stress granule marker. Cytoplasmic droplet formation for hSSB1-EGFP upon oxidative stress is even more pronounced than that for endogenous hSSB1; however, hSSB1-EGFP droplets appeared even in the absence of $H_2O_2$ treatment. Even untreated cells show cytoplasmic granulation of G3BP1, which was not seen in untransfected cells **(**cf. **Fig. 6A)**. Cytoplasmic condensates, however, were only visible in red channel (G3BP1 signal) and not in green (EGFP) when EGFP alone was overexpressed. **(B)** Top row shows confocal images of a fixed HeLa cell overexpressing hSSB1-EGFP treated with 0.3 mM $H_2O_2$. hSSB1-EGFP and G3BP1 show colocalized enrichment in cytoplasmic droplets. Green channel shows hSSB1-EGFP, red channel shows G3BP1. Yellow color indicates colocalization. Bottom row shows 3D reconstructed images of the same ROI from confocal image Z-stacks. **(C)** Cell typization data showing similar fractions of SG+ cells upon both EGFP and hSSB1-EGFP overexpression with no significant differences, indicating that hSSB1 overexpression *per se* does not influence SG formation. Means ± SEM are shown together with individual data points (two-sample T-tests, $p < 0.05$). Ns, not significant.

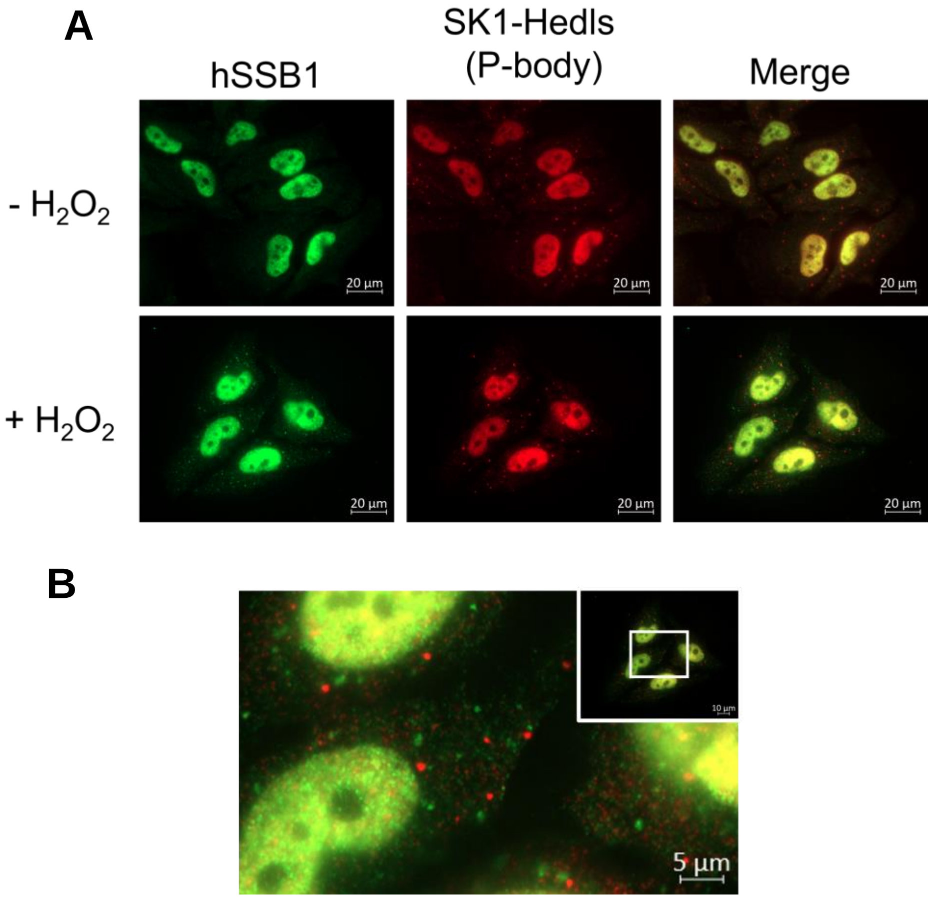


**Fig S15. Cytoplasmic hSSB1 granules do not colocalize with P-bodies**.
(**A**) Representative epifluorescence images (*n* = 3 independent measurements) of untreated and $H_2O_2$-treated (300 µM, 2 h) HeLa cells. Green and red channels show endogenous hSSB1 and SK1-Hedls P-body marker, respectively. P-bodies were visible even in untreated cells, while hSSB1 granules formed only under oxidative stress conditions. (**B**) Magnified epifluorescence image of a $H_2O_2$-treated HeLa cell from panel **A**. No colocalization was observed between hSSB1 granules and P-bodies.

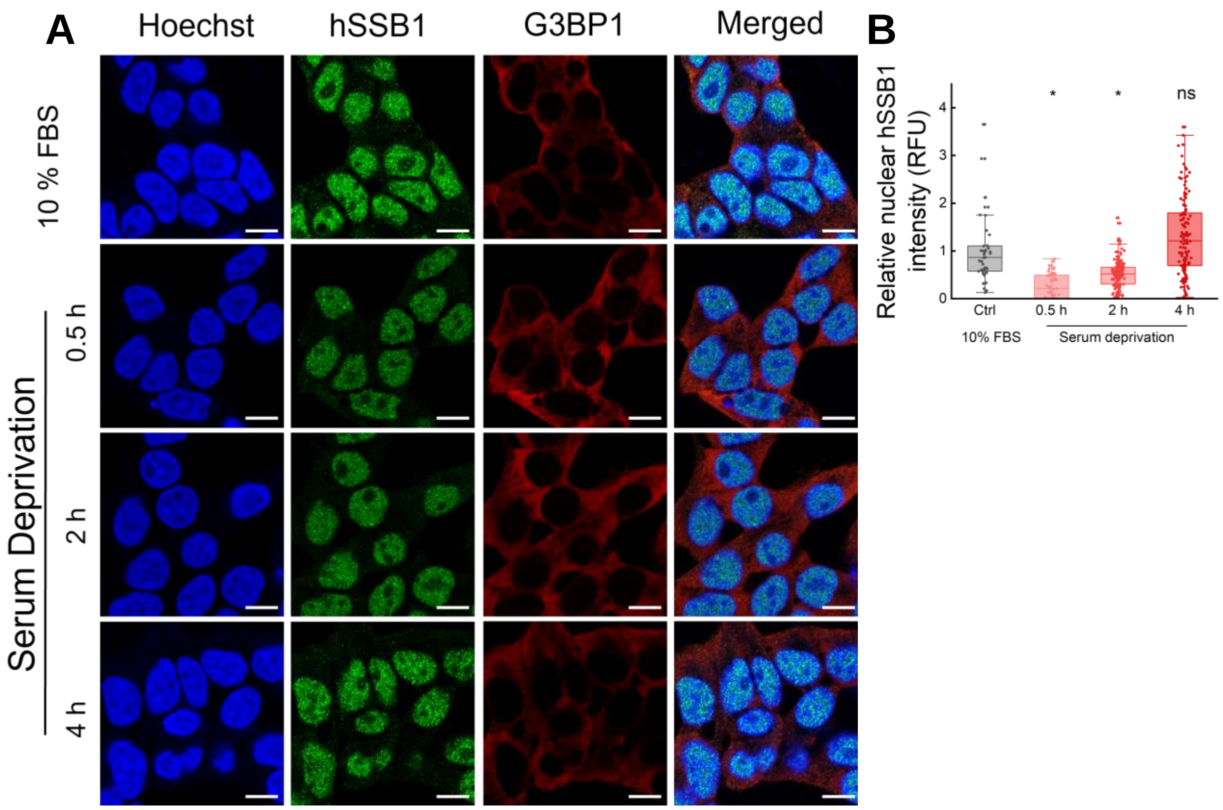


**Fig S16. Serum deprivation causes a rapid decrease in nuclear hSSB1 intensity that is restored over time**

(**A**) Representative confocal microscopic images of immunostained HeLa cells that were kept in serum-free media for the indicated times. Blue channel shows nuclear Hoechst stain; green and red channels show endogenous hSSB1 and G3BP1 stress granule marker, respectively. Merged image was created from all three channels. Scale bar: 10 µm. (**B**) Serum deprivation results in a rapid (0.5 h) decrease in nuclear hSSB1 signal, which is restored after 4 h. Relative nuclear hSSB1 intensities (RFU, relative fluorescence units) were normalized to those in serum-containing control. Dots indicate individual nuclear intensities (Kruskal-Wallis-test with Dunn's post-hoc test, * indicates significant difference, 'ns' not significant, $p < 0.05$).

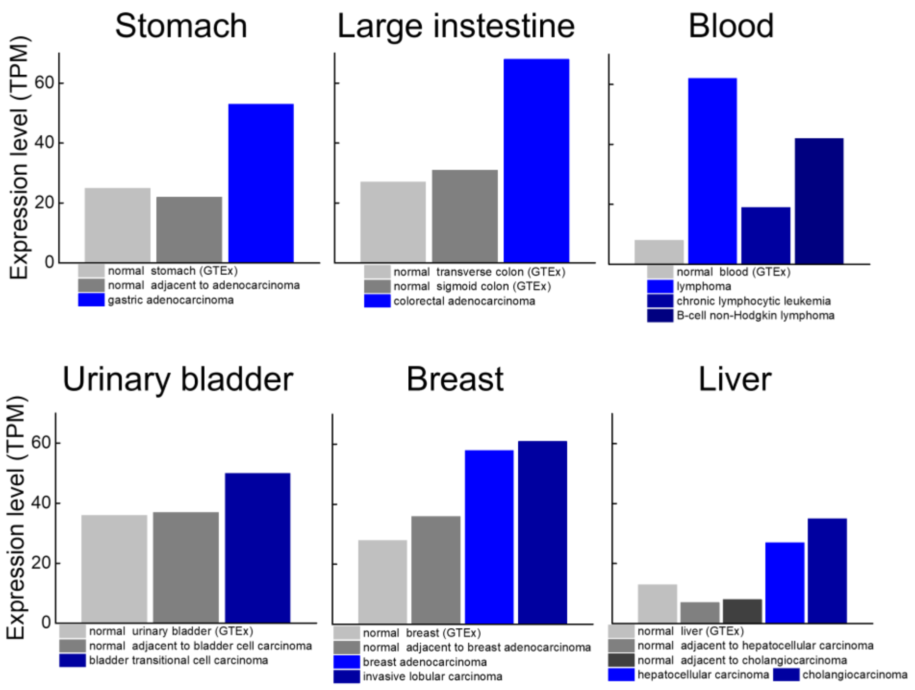


**Fig. S17. hSSB1 shows elevated expression levels in various cancerous tissues**
hSSB1 expression levels, displayed as TPM (transcripts per million), are shown for non-cancerous (grey shades) and cancerous (blue shades) tissues (GTEx: Genotype-Tissue Expression, data source: Expression Atlas, https://www.ebi.ac.uk/gxa/home).

### Supplementary Tables

| **Protein** | **Substrate** | ***K*d (µM)** | ***n*** |
|---|---|---|---|
| hSSB1 (WT) | ssDNA | 0.041 ± 0.002 | 1.7 ± 0.1 |
| hSSB1 (WT) | ssRNA | 0.054 ± 0.004 | 1.5 ± 0.2 |
| hSSB1-C41S | ssDNA | 0.042 ± 0.006 | 1.4 ± 0.3 |
| hSSB1-C81S | ssDNA | 0.029 ± 0.003 | 1.7 ± 0.3 |
| hSSB1-C99S | ssDNA | 0.015 ± 0.003 | 1.6 ± 0.4 |
| hSSB1-C41S-C81S | ssDNA | 0.030 ± 0.005 | 1.3 ± 0.3 |
| hSSB1-C41S-C99S | ssDNA | 0.043 ± 0.006 | 1.8 ± 0.4 |
| hSSB1-C81S-C99S | ssDNA | 0.026 ± 0.004 | 1.6 ± 0.4 |
| hSSB1-C41S-C81S-C99S | ssDNA | 0.028 ± 0.002 | 1.7 ± 0.2 |
| hSSB1-dIDL | ssDNA | 0.68 ± 0.05 | 1.7 ± 0.2 |

**Table S1. ssDNA and ssRNA binding parameters of hSSB1 variants determined in fluorescence anisotropy titrations.** Table shows determined equilibrium dissociation constants ($K_d$) and Hill coefficients ($n$) together with their fitting error (see Materials and Methods for further details).
